# Comparative connectomics of two distantly related nematode species reveals patterns of nervous system evolution

**DOI:** 10.1101/2024.06.13.598904

**Authors:** Steven J. Cook, Cristine A. Kalinski, Curtis M. Loer, Nadin Memar, Maryam Majeed, Sarah Rebecca Stephen, Daniel J. Bumbarger, Metta Riebesell, Barbara Conradt, Ralf Schnabel, Ralf J. Sommer, Oliver Hobert

## Abstract

Understanding the evolution of the bilaterian brain requires a detailed exploration of the precise nature of cellular and subcellular differences between related brains. To define the anatomical substrates of evolutionary change in the nervous system, we undertook an electron micrographic reconstruction of the brain of the predatory nematode *Pristionchus pacificus.* A comparison with the brain of *Caenorhabditis elegans*, which diverged at least 100 million years ago, reveals a conserved nematode core connectome and a wide range of specific substrates of evolutionary change. These changes include differences in neuronal cell death, neuronal cell position, axo-dendritic projection patterns and many changes in synaptic connectivity of homologous neurons that display no obvious changes in overall neurite morphology and projection patterns. Arguing against specific hot spots of evolutionary change, connectivity differences are distributed throughout the nervous system and extend to glia as well. We observed examples of apparent circuit drift, where changes in morphology and connectivity of a neuron do not appear to alter its behavioral output. In conclusion, our comprehensive comparison of distantly related nematode species provides novel vistas on patterns of conservation as well as the substrates of evolutionary change in the brain that span multiple organizational levels.

---

Brains have undergone dramatic changes across animal evolution, evidenced by elaborations and increases in size, anatomical complexity and functional capacities. Due to anatomical complexity and difficulties associated with identifying homologous neurons, the precise natures of evolutionary changes are often hard to pinpoint with cellular or synaptic resolution in vertebrate brains. However, in organisms with numerically more constrained and anatomically well-described brains, identifying homologous neurons is more feasible, which permits the delineation of the precise substrates of anatomical change in the brain. Such analysis can address several key questions (Roberts *et al*. 2022): What are the exact neuroanatomical substrates of evolutionary change (e.g., neuron gain or loss; changes in neuronal soma migration patterns; changes in projection patterns; synaptic connectivity changes)? Do some types of changes happen more often than others? Are specific types of changes homogenously distributed across the network or are there hotspots of evolutionary changes (e.g., is the sensory periphery of an animal more evolvable than central circuits)?

The nematodes *Pristionchus pacificus* and *Caenorhabditis elegans* are thought to have diverged more than 100 million years ago (Qing *et al*. 2023). While this substantial divergence is not immediately obvious based on the overall size or morphology of these species, it shows on many other levels. On a molecular level, the *P. pacificus* genome is 70% larger than the *C. elegans* genome, contains 30% more genes and only roughly 30% of its genes have 1:1 orthologs in *C. elegans* (Athanasouli et al. 2020). The divergence of these two nematode species is also apparent in their adaptation to distinct ecological niches and in unique adaptions in sensory, locomotory and feeding behavior (Hong and Sommer 2006; Srinivasan *et al*. 2008; Kroetz *et al*. 2012). The most striking of these adaptations is the predatory feeding behavior that *P. pacificus* displays toward other nematodes and an associated self-recognition system that prevents cannibalism against kin (Ragsdale *et al*. 2013; Lightfoot *et al*. 2019).

The many cataloged differences between *P. pacificus* and *C. elegans,* particularly on the behavioral level, prompt the question to what extent the nervous system of these two species has diverged. Previous studies using serial section electron microscopy described differences in the sensory periphery of these worms (i.e., the exposed ciliated ending of these two species) and the small and isolated nervous system of the foregut (“pharynx”) (Bumbarger *et al*. 2013; Hong *et al*. 2019). We analyzed to what extent the entire centralized brain - the cluster of head ganglia that controls many complex nematode behaviors – has diverged among these two distant nematode species. We expected that such analysis might identify and provide a panoramic view of the substrates of evolutionary change, thereby addressing several questions that only a broad, brain-wide comparative connectomic analysis can answer.

## Nanoscale reconstruction of the *P. pacificus* nervous system

We analyzed serial section electron microscopy of two adult hermaphrodite *P. pacificus* heads from the anterior nose tip to the retrovesicular ganglion, including the nerve ring, the major neuropil of nematodes (**Figure 1A**). We reconstructed the ultrastructure of all neurons in these two animals, analyzed the extent of the membrane contacts between their neurites (the “contactome”) (**Supplement Data 1,2**) and their patterns of chemical synaptic connectivity (“connectome”) **(Supplement Data 3,4**). Based on remarkably well-conserved cell body position as well as characteristic features of neurite trajectories and placement within fascicles, we were able to assign homology to all neurons found in the *C. elegans* head ganglia (**Figure 1B; Figure 1 – Supplement 1;** renderings of all neurons are shown in **Supplemental File 1**). Such clear one-to-one homology assignments are difficult to make upon comparison of more complex nervous systems and present a unique advantage of the nematode phylum with its limited cellular complexity and apparent stereotypy of cellular composition and lineage. These neuronal homology assignments allowed us to ask if - and to what extent - homologous neurons have evolved distinct features. In the ensuing sections of this manuscript, we describe distinct patterns of evolutionary changes observed.

**Figure 1:**
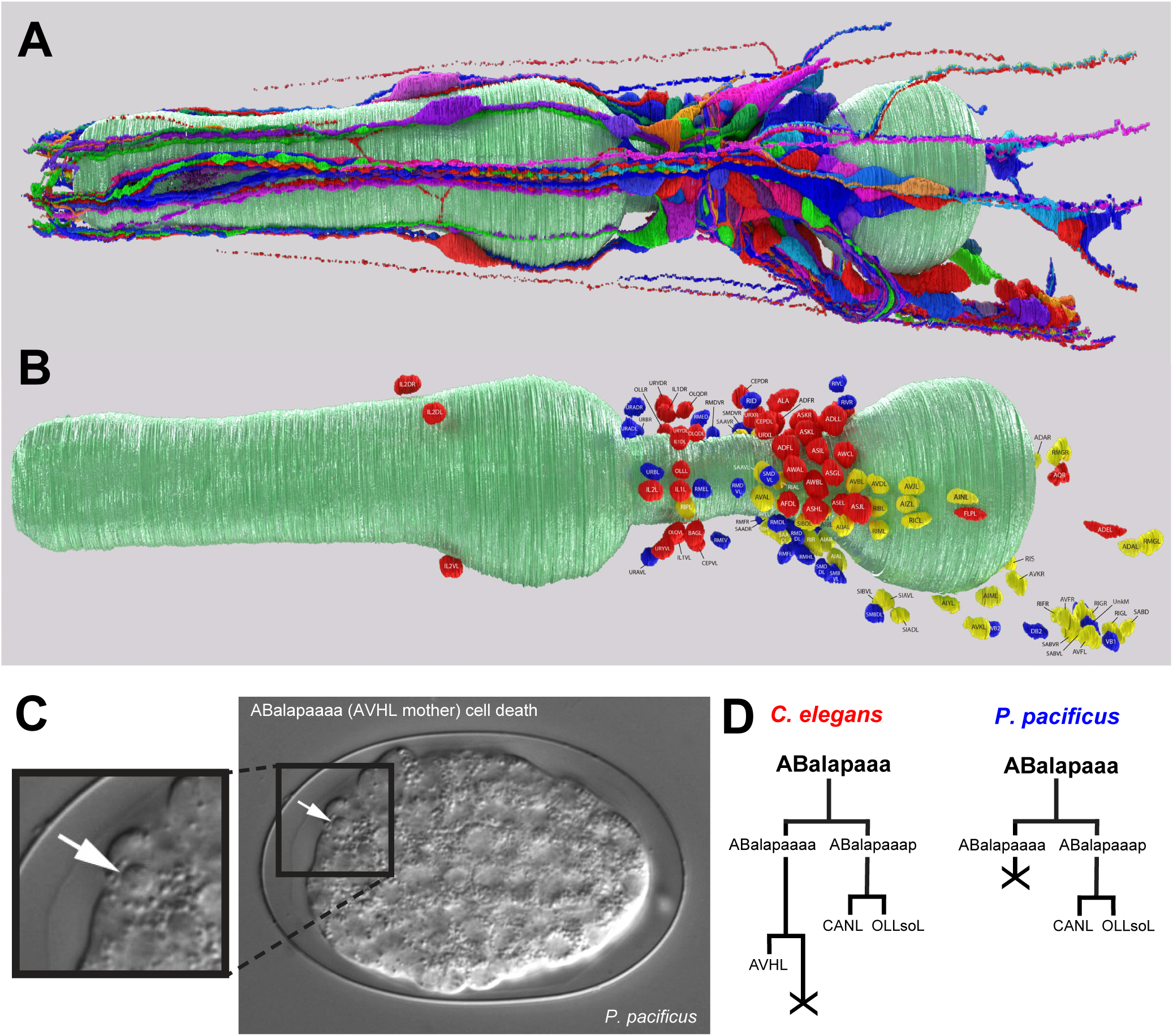
The reconstructed *Pristionchus pacificus* brain. **A:** A left lateral view 3D rendering of all *P. pacificus* neurons reconstructed from serial section electron micrographs of the head. Neurons are shown in different colors for contrast, the pharynx is shown in teal. **B:** A Left lateral view 3D rendering of all neuronal nuclei in the *P. pacificus* head. Nuclei are colored by neuronal function experimentally determined in nematodes: Sensory – red, interneuron – yellow, motor – blue, the pharynx is shown in teal. See **Figure 1 Supplement 1** for different perspective of neuronal nuclei position) **C:** Differential interference contrast image of *P. pacificus* embryo, indicating dying progenitor cell responsible for AVH neuron loss in *P. pacificus* compared to *C. elegans*. See **Figure 1 Supplement 2A** for detailed lineage data. **D:** Lineaging diagram displaying the death of AVH’s sister cell in *C. elegans* (left) and AVH’s progenitor cell death in *P. pacificus* (right) during embryogenesis. See **Figure 1 Supplement 2A** for detailed lineage data.

## Differences in neuronal composition and location

We found one *C. elegans* interneuron class, the bilaterally symmetric AVH neuron pair, to be missing from the *P. pacificus* head region. This neuron pair is stereotypically located in the lateral ganglion in *C. elegans,* from where it extends a neurite through the nerve ring into and along the ventral nerve cord (White *et al*. 1986). The sister cells of both left and right AVH neurons undergo programmed cell death in *C. elegans* (Sulston et al. 1983)(**Figure 1C, Figure 1 Supplement 2A**). Tracing the lineage of embryonic cell division in *P. pacificus* using 4D video microscopy (Schnabel et al. 1997), we found that this cell death occurs one cell division earlier compared to *C. elegans*, hence explaining the absence of AVHL/R **(Figure 1C, D; Figure 1 Supplement 2B**). Our embryonic lineaging analysis also reveals that precocious cell death can explain the previously noted absence of the pharyngeal gland cell g2 in *P. pacificus* (Riebesell and Sommer 2017). As in the case of AVH, the sister of g2 dies in *C. elegans*, while in *P. pacificus,* the mother dies (**Figure 1 Supplement 3A-D**). Hence, together with the previously reported excess cell death of a subset of the VC ventral cord motorneuron class in *P. pacificus* (Eizinger et al. 1999), alterations in the spatial and temporal specificity of programmed cell death appear to constitute a commonly used mechanism for shaping species-specific nervous system composition.

We traced the lineage of AVH in two more distantly related nematodes that can serve as outgroups to *P. pacificus* and *C. elegans*, *Acrobeloides nanus* and *Plectus sambesii,* revealing that in both species the cleavage pattern of the AVH-generating lineage (as well as the g2 gland-generating lineage) resembles the *C. elegans* pattern. In both species, the sister, rather than the mother, of the presumptive AVH neuron and g2 gland cell undergoes programmed cell death (**Figure 1 Supplement 4A-C**). These observations indicate that the loss of *P. pacificus* AVH and other cells due to the heterochronic death of the mother cell is a repeated, derived feature of embryogenesis in *P. pacificus*.

We observed a remarkable molecular correlation to the loss of AVH neurons in *P. pacificus.* In *C. elegans*, the AVH neurons require a divergent member of the bHLH-PAS family of transcription factors, *hlh-34*, for their proper differentiation (Berghoff *et al*. 2021; Cook *et al*. 2021). Sequence surveys of multiple highly divergent nematode species with complete and well-annotated genomes show that the *hlh-34* locus is lost in the *Pristionchus* genus (**Figure 1 Supplement 2C**). Hence, the loss of a neuron due to heterochronic cell death is correlated with the genomic loss of its identity regulator.

We asked whether the loss of AVH in *P. pacificus* is accompanied by synaptic wiring changes in a neighboring neuron, AVJ, that shares many anatomical similarities to AVH in *C. elegans,* with both neurons displaying a similar neurite extension pattern through the nerve ring and along the ventral cord (White *et al*. 1986). The two neurons are electrically coupled and share some synaptic partners (White *et al*. 1986; Cook *et al*. 2019; Witvliet *et al*. 2021). However, we do not observe that the synaptic contacts normally specifically made by AVH and not AVJ are absorbed by *P. pacificus* AVJ. Instead, AVJ generates additional novel, *P. pacificus-*specific synapses (see diagrams below).

## Differences in neuronal location

Another type of evolutionary change concerns cell soma position. Generally, the relative positions of homologous neuronal somas are highly conserved between *P. pacificus* and *C. elegans*. Notable exceptions are the inner labial, dorsal and ventral IL2 (IL2D and IL2V) sensory neuron pairs. In *C. elegans,* the IL2D and IL2V neuronal soma are located at a similar position along the longitudinal axis in the anterior ganglion as the lateral IL2 neuron pair (White *et al*. 1986). In *P. pacificus,* however, the IL2D and IL2V soma pairs are positioned much more anteriorly (**Figure 1B**). Their axons therefore need to take a much longer path to reach the nerve ring, a feat that they achieve by close fasciculation with inner labial IL1 neurons, whose position is unchanged relative to *C. elegans* (**Figure 1 Supplement 5**). The more anterior localization of the IL2D/V soma results in substantially shortened dendritic length. Since the *C. elegans* IL2D/V dendritic generate branches during the dauer stage to respond to mechanosensory touch involved in nictation behavior (Lee *et al*. 2012; Schroeder *et al*. 2013), such shortening may result in a reduced sensory perceptive field of the branched IL2D/V dendrites in nictating *P. pacificus* dauer stage animals.

## Differences in neurite projection patterns alter network architecture

Most individual neurons display highly characteristic neurite projection or branching patterns in *C. elegans,* which we leveraged to describe similarities and differences across species. For example, the *C. elegans* AIB neurites display an abrupt neighborhood shift in the nerve ring (White *et al*. 1986) which is conserved in *P. pacificus* (**Supplemental File 1**)(Hong *et al*. 2019). Similarly, the unusual, truncated dendrite of the amphid-associated *C. elegans* AUA neuron class (White *et al*. 1986) is also conserved in *P. pacificus* (**Supplemental File 1**). The ring interneurons RIM, RIH, RIS and RIR display characteristic branching patterns in the *C. elegans* nerve ring and these patterns are also similar in both species (**Supplemental File 1**). Paralleling these notable patterns of conservation, we discovered a rich set of evolutionary diversifications of projection patterns; several of them resulting in striking differences in presumptive information flow in the two nervous systems. Among the most dramatic differences is a complete repurposing of adult URB neuron function. In *C. elegans*, all available EM constructions, as well as reporter genes, show that URB extends a long dendrite toward the tip of the nose along the lateral amphid sensory dendrite bundle and extends an axon into the nerve ring to synapse with distinct sensory, inter- and motorneuron classes (White *et al*. 1986; Cook *et al*. 2019; Witvliet *et al*. 2021). In striking contrast, no dendritic extension of URB neurons is observed in either of the two *P. pacificus* samples (**Figure 2A, Figure 1 Supplement 1**). Moreover, unlike in *C. elegans,* the *P. pacificus* URB axon innervates the head muscles in adults (**Figure 2 Supplement 1A**). The concordant loss of dendrite and rewiring of axonal output suggests a functional switch from sensory to motorneuron between these two species.

**Figure 2:**
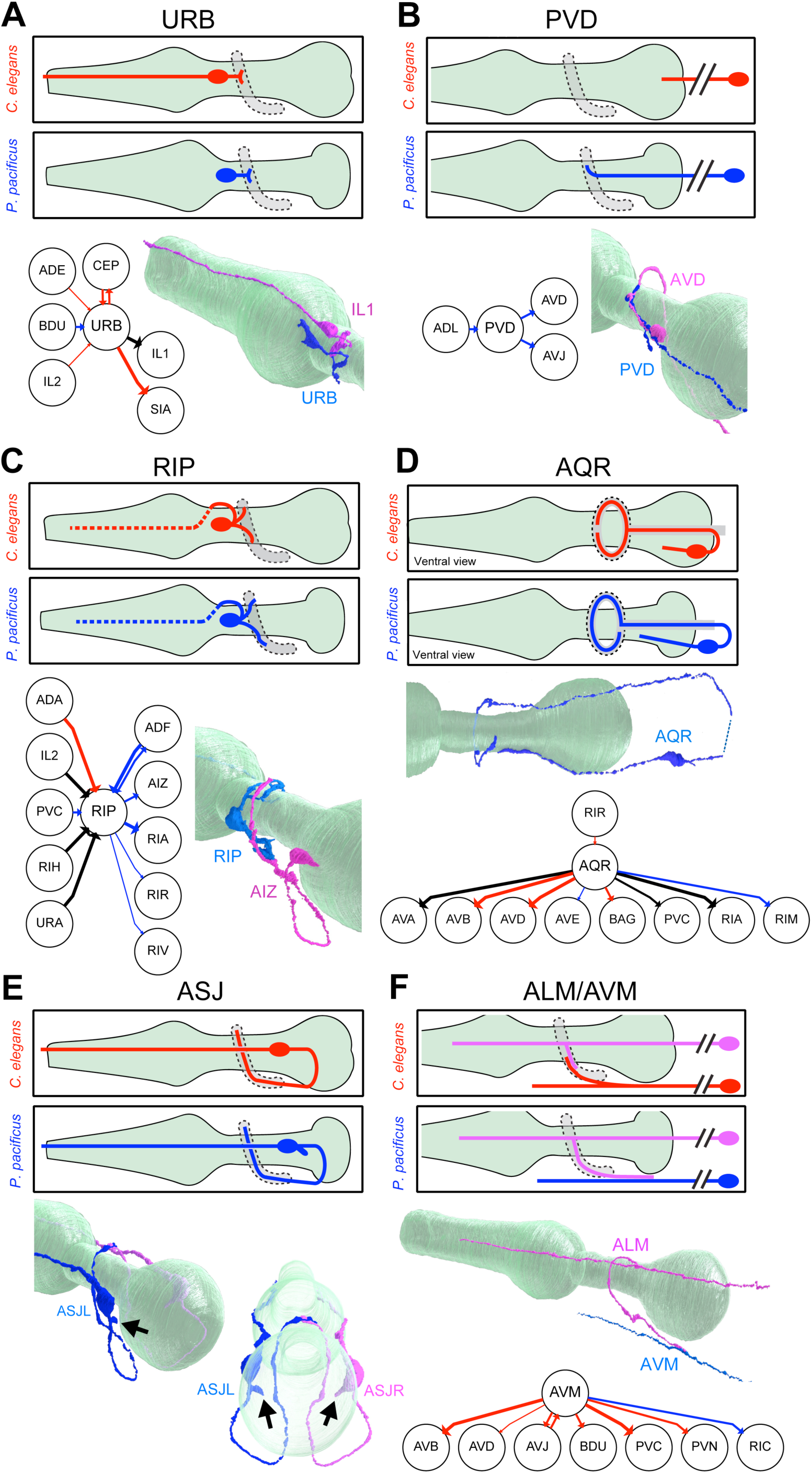
Morphological evolution of neurite trajectories. In each panel (**A-E**), the top section shows a schematic comparison of the neuron’s morphology in *C. elegans* (red) and *P. pacificus* (blue). Schematics are left lateral views (anterior to the left and dorsal up) for all panels except **C** (AQR), which is a ventral view. The pharynx is shown in green and the nerve ring (neuropil region) is grey, outlined by a dashed line. Below the schematics are a connectivity diagram and 3D rendering of the *P. pacificus* neuron (blue), in some cases with a partner neuron (magenta). Connectivity diagrams show chemical synapses that are conserved in all adult nematode samples (black), are *C. elegans*-specific (red), or *P. pacificus*-specific (blue). Line width is proportional to synaptic weight and arrowheads show directionality. (A connectivity diagram is not relevant for panel E, and it not included.) **A:** URB morphology, illustrating the absence of a dendrite in *P. pacificus.* The 3D rendering of *P. pacificus* URB (blue) is shown with a conserved synaptic partner of URB, the inner labial sensory neuron IL1 (magenta), which retains its dendrite in *P. pacificus*, extending from the nerve ring region to beyond the anterior tip of the pharynx. **B:** PVD morphology, illustrating that in *P. pacificus* PVD extends into the nerve ring where it synapses with AVJ and AVD (seen in both *P. pacificus* samples). The double slash indicates the PVD soma is posterior to the reconstruction. The 3D rendering shows PVD (blue) with *P. pacificus*-specific synaptic partner AVD (magenta), a closeup of the nerve ring region. **C:** RIP morphology, illustrating extended nerve ring neurites in *P. pacificus.* The 3D rendering shows RIP (blue) r with *P. pacificus*-specific synaptic partner AIZ (magenta), a closeup of the nerve ring region. **D:** AQR morphology *(ventral view),* illustrating how the *P. pacificus* AQR neuron crosses over the dorsal midline rather than bifurcating ventrally as seen in *C. elegans*. The 3D rendering shows AQR (blue) entering the nerve ring on the left (top), extending over dorsally to the right side (bottom). The AQR dendrite extends along the right side (bottom) almost to the nerve ring. The dashed line indicates that the posterior-most AQR connecting neurite is not included in the reconstruction. **E.** ASJ morphology, illustrating branches protruding from or near the ASJ soma in *P. pacificu*s not found in *C. elegans*. 3D renderings of both ASJL (blue) and ASJR (magenta) are shown in the lateral posterior view (left) and dorsal posterior view (right, with nearly transparent pharynx), showing that these branches, indicated by arrows, project near the pseudocoelom. **F.** ALM and AVM morphologies illustrate that in *P. pacificus,* AVM does not send a neurite into the nerve ring (whereas in *C. elegans* it forms a gap junction with ALM). Instead, ALM extends a neurite to meet ALM ventrally, where a gap junction is formed. The double slash indicates the AVM and ALM somas are posterior to the reconstruction. The 3D rendering shows the simpler *P. pacificus* AVM neurite (blue) contacted by an extended ALM neurite (magenta). The connectivity diagram shows many *C. elegans*-specific synapses made in the nerve ring by AVM, which no longer contacts these cells in *P. pacificus*.

Altered neurite extension patterns within the lateral nerve also indicate a divergence of routes of synaptic communication in the two nematode species. In *C. elegans,* the PVD sensory neuron extends a primary dendrite that typically terminates before reaching the nerve ring (White *et al*. 1986; Cook *et al*. 2019). In marked contrast, in both *P. pacificus* samples, this dendrite projects further anteriorly into the nerve ring where it synapses onto AVD and AVJ and receives inputs from the nociceptive ADL neuron (**Figure 2B, Figure 2 Supplement 1B**). Hence, the PVD neurite, normally purely dendritic in *C. elegans,* is repurposed to become axodendritic in *P. pacificus*, thus allowing the lateral nerve to provide ascending input to the brain.

Conversely, the harsh touch mechanoreceptive neuron, FLP, has a shorter axon in *P. pacificus*. In *C. elegans,* the two FLP neurons extend their axons through the deirid commissure, then project into the nerve ring where they generate synaptic outputs to the pre-motor command interneuron AVA (White *et al*. 1986; Cook *et al*. 2019; Witvliet *et al*. 2021). In *P. pacificus,* the FLP axons terminate before the nerve ring resulting in a loss of connections to the backward movement-inducing AVA. Meanwhile, new connections to the forward movement-inducing PVC command interneuron are seen (**Figure 1 Supplement 3**). While the precise function of FLP in *P. pacificus* is unknown, altering its output from AVA to PVC suggests a repurposing from avoidance to attractive responses to head touch. This change may relate to the predatory behavior of *P. pacificus*, in which contact with prey may not elicit an escape response.

A striking alteration in information flow is apparent in the RIP neuron pair, the only neuron that synaptically connects the somatic and pharyngeal nervous systems (White *et al*. 1986; Bumbarger *et al*. 2013; Cook *et al*. 2019; Witvliet *et al*. 2021). At all developmental stages, the posteriorly directed neurite of RIP is strictly dendritic in the nerve ring of *C. elegans*, assembling synaptic input from a variety of central neuron classes (Witvliet *et al*. 2021). In *P. pacificus,* the posteriorly directed RIP neurite extends more deeply into the nerve ring and generates synaptic outputs onto different neuron types (**Figure 2C**). Hence, synaptic information flow through RIP appears unidirectional in *C. elegans* (from the nerve ring to the enteric nervous system), while in *P. pacificus*, RIP undergoes a partial polarity change to directly synapse onto both the somatic and enteric nervous systems. It will be interesting to see whether this rewiring relates to the predatory feeding behavior displayed by *P. pacificus*.

The unpaired AQR neuron, an oxygen-sensing neuron exposed to internal body cavities in both species, undergoes another example of striking projection change. In *C. elegans,* the AQR neurite bifurcates upon entry into the nerve ring to extend symmetric neurites along either side of the nerve ring which then terminates shortly before reaching the dorsal midline (**Figure 2D**)(White *et al*. 1986; Witvliet *et al*. 2021). In *P. pacificus*, the neurite of AQR does not bifurcate; instead, a single neurite reaches across the dorsal midline to the other side of the nerve ring (**Figure 2D**). The *P. pacificus* AQR neuron makes three small branches not observed in *C. elegans* as well as a large swelling at the dorsal midline, resulting in a preservation of distinctive gap junctions to the PVP neuron, as observed in *C. elegans* (White *et al*. 1986)**(Supplemental File 1**). However, the *P. pacificus* AQR neuron does not innervate pre-motor command interneurons AVB and AVD, suggesting that AQR activation may trigger distinct behavioral read-outs. Indeed, oxygen-induced behavioral responses are divergent between *P. pacificus* and *C. elegans* (Moreno *et al*. 2016).

The ASJ amphid sensory neurons display another type of projection change. ASJ neurons in both *C. elegans* and *P. pacificus* display a bipolar morphology with a sensory dendrite to the tip of the nose and an axon into the nerve ring. In *P. pacificus*, both ASJ neurons extend an additional short, thick neurite at or near its soma, not directed toward a specific fascicle but extending along amphid cell bodies **(Figure 2E)**. Strikingly, in both samples and on both sides of the animal, this short neurite is filled with dense core vesicles (**Figure 2 Supplement 1C**), indicating that in *P. pacificus* this neuron has the additional capacity to broadcast neuropeptidergic signals across the nervous system.

Changes in projection patterns do not necessarily result in changes in information flow. The two anterior light touch receptor neuron classes ALM and AVM are electrically coupled, and each innervates command interneurons to signal reversal behavior in *C. elegans* (Chalfie *et al*. 1985; White *et al*. 1986). The ventrally located AVM projects a bifurcating axon into the *C. elegans* nerve ring, where synapses are made to command interneurons, and terminates after meeting branches that are sent by lateral ALM touch sensory neurons into the nerve ring (**Figure 2F**)(White *et al*. 1986; Witvliet *et al*. 2021). In marked contrast, the AVM neuron does not send a bifurcating branch into the *P. pacificus* nerve ring (**Figure 2F**), resulting in the loss of many AVM synapses observed in the adult *C. elegans*. However, the *P. pacificus* ALM neurites instead project further ventrally and posteriorly in *P. pacificus* to reach the unbranched AVM neurite on the ventral side (**Figure 2F**), generating the same large gap junction connection to AVM as they do in *C. elegans* (**Figure 2 Supplement 1D**). In aggregate, synaptic outputs of the ALM and AVM neuron classes to command interneuron are therefore preserved in both nematodes, which is manifested by a similar gentle touch response in both species.

## Synaptic connectivity repatterning without obvious changes in neurite projection

We discovered extensive evolutionary changes in synaptic connectivity patterns in the nerve ring independent of any obvious changes in overall neuron morphology. Analysis of multiple *C. elegans* connectomes across development has indicated a large degree of variability in synaptic connectivity patterns (Witvliet *et al*. 2021). To account for such variability in the context of interspecies comparisons, we first extracted all synaptic connections that are completely conserved in all 10 available *C. elegans* samples covering multiple developmental stages (L1, L2, L3, L4, Adult) and our two adult *P. pacificus* samples. These connections define a “core connectome” that is conserved across development and evolution. In addition, we identified connections that are conserved in both *P. pacificus* samples but absent from all 10 *C. elegans* samples (*P. pacificus*-specific connectome) and connections that are absent in both *P. pacificus* samples but present in all 10 *C. elegans* samples (*C. elegans-*specific connectome). These sets of synaptic classifications are illustrated in (**Figure 3A-D)**. Schematic connectivity diagrams of each neuron class are assembled in **Supplemental File 2.**

**Figure 3:**
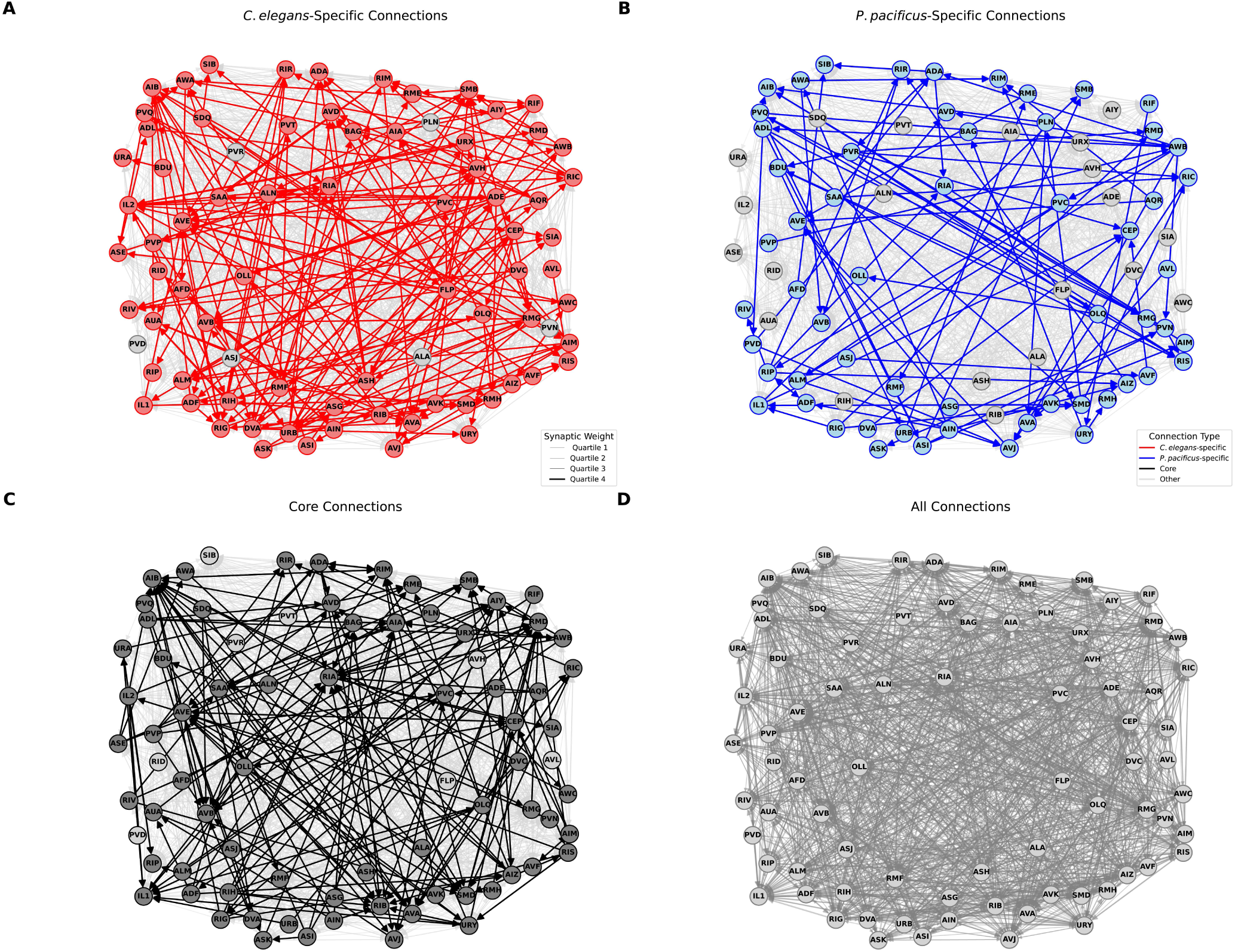
Connectomic comparison reveals widespread connectivity changes. Panels A-D: Network (wiring) diagrams showing all neurons in the nerve ring and their synaptic connections: core (black), variable (gray), *P. pacificus*-specific (blue), *C. elegans*-specific (red). Arrow widths are proportional to the average number of synapses across datasets; arrowheads represent directionality. **A**. Diagram highlighting *C. elegans*-specific connections in red. Legend for synaptic edge weights is provided. **B.** Diagram highlighting *P. pacificus*-specific connections in blue. Legend for color code is provided. **C:** Core wiring diagram highlighting shared, non-variable connections in black. **D.** Complete wiring diagram showing all *P. pacificus* and *C. elegans* connections.

We find that 88% of all neuron classes contribute to the “core nematode connectome” observed in every available EM sample from both species, independent of developmental stage. The core connectome is not biased to a specific part of the nervous system, i.e., it includes sensory, inter- and motorneurons and extends from the sensory periphery via interneuronal layers to various motor outputs (**Figure 3**). Conversely, neurons that do not form any “core nematode connections” (AVH, AVL, CAN, FLP, PVR, PVT, RID, and SIB) fall into multiple functional categories and are also not biased to a specific part of the nervous system.

Species-specific synaptic connections are also not limited to a specific part of the nervous system, such as its sensory periphery, but are rather very pervasively distributed throughout the entire connectome. In total, 96% of the analyzed neurons of the adult *P. pacificus* and *C. elegans* brain make species-specific connections, 72% of neurons generate *P. pacificus*-specific connections, while 87% of neurons generate synaptic partnerships only in *C. elegans* (**Figure 3; Figure 3 Supplement 1**). Differences in the percentage of species-specific synapses are likely a reflection of a smaller sample size in *P. pacificus* compared to *C. elegans*. Species-specific synapses account, on average, for only 10.87% and 4.92% of the synaptic connections of each neuron in *C. elegans* and *P. pacificus,* respectively. (**Figure 3 Supplement 1**). Only three neuron classes, ALA, CAN and SAB do not form species-specific synapses (**Figure 3C, D, Supplemental File 2**). Other signaling capacities of the ALA, e.g., by the dense core vesicle-filled *P. pacificus-*specific extension of ASJ (see above), may have diverged between these two species, while the primary function of CAN is likely neuropeptidergic (Ripoll-Sanchez *et al*. 2023).

## Synaptic wiring changes as a function of neighborhood changes

Species-specific synapses of morphologically similar neurons can be generated by dramatic shifts in the neuronal neighborhood. The most striking example is the polymodal, nociceptive ASH neuron. In *C. elegans,* the axon of ASH extends medially through the nerve ring, making ample *en passant* contacts with several interneurons, most notably command interneurons, until it meets its contralateral homolog at the dorsal midline (**Figure 4A, Supplement 1**). The synaptic output of ASH transforms ASH-sensed nociceptive stimuli into a reversal response via pre-motor command interneurons (Kaplan and Horvitz 1993). Like in *C. elegans,* each ASH axon in *P. pacificus* also extends through the nerve ring to meet its contralateral homolog at the midline, but the path taken through the nerve ring is dramatically different as it mostly travels along the outer side of the nerve ring (**Figure 4B**). This dramatic difference in the neuronal neighborhood has substantial consequences on the *en passant* synaptic connectivity: ASH loses its direct output to the homologs of the backward locomotion-inducing command interneurons in *P. pacificus* (**Figure 4C**). These differences in synaptic connectivity are particularly remarkable in light of the experimental finding that in *P. pacificus,* the ASH neurons also mediate reversal to nociceptive stimuli as in *C. elegans* (Srinivasan et al. 2008). Hence, in these two nematodes, nociceptive sensory information is perceived by the same neuron but appears to be relayed through different synaptic pathways to produce similar motor outputs.

**Figure 4:**
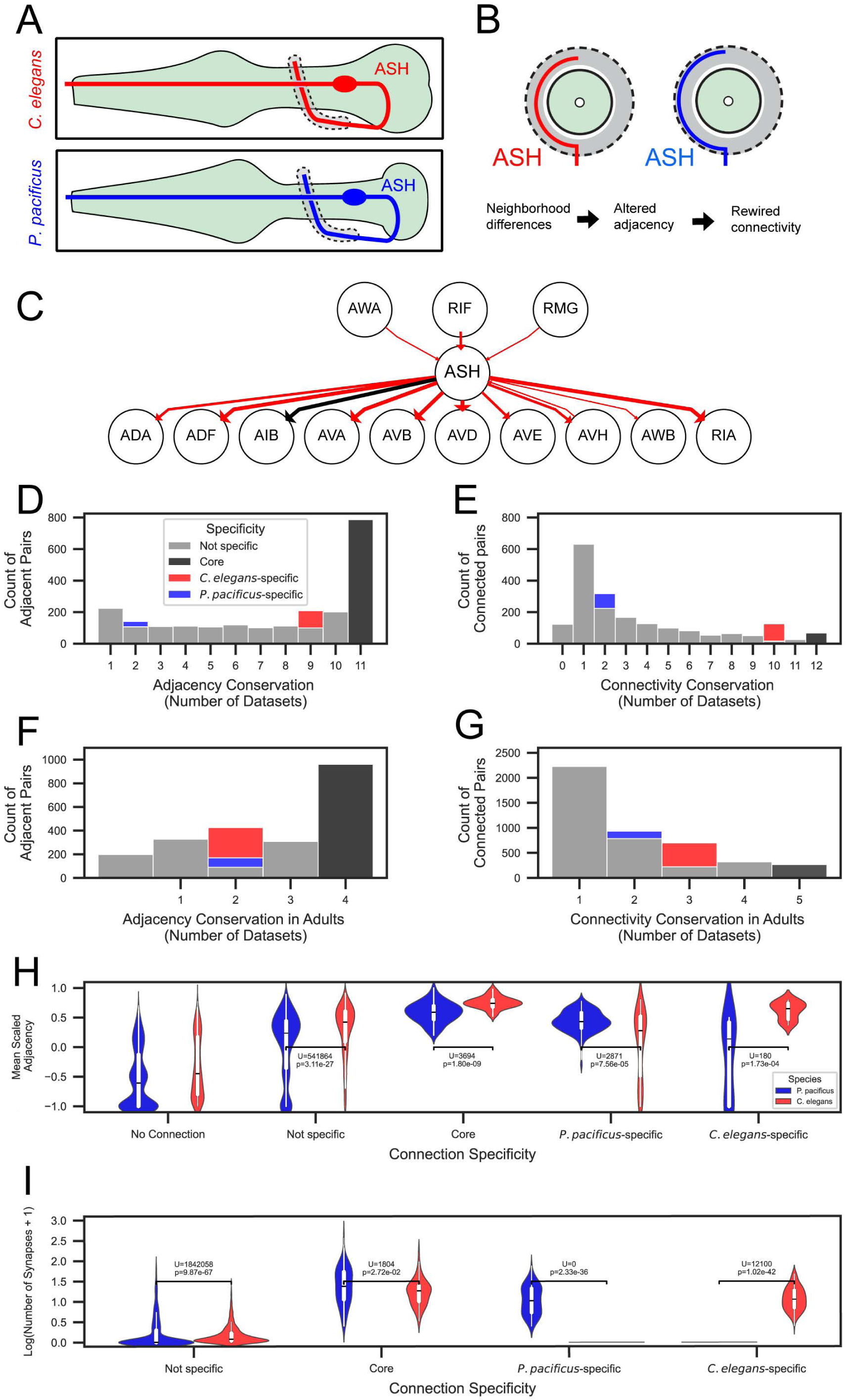
Neurite neighborhood shifts and quantitative analysis of synaptic connectivity differences. **A:** Schematic comparison of ASH neuron morphology in *C. elegans (red)* and *P. pacificus (blue),* showing similar morphology at a gross anatomical scale. Left lateral view (see legend for Fig 2). **B:** Schematic of nerve ring cross section showing lateral displacement of ASH neurite in *P. pacificus* (blue) compared to *C. elegans* (red). The shift in neighborhood is shown in a diagram as a driver of synaptic rewiring. **C:** Adult wiring diagram of ASH. *C. elegans*-specific connections are shown in red. **D:** Histogram of adjacency conservation (Number of EM series with edge present) among all nematode nerve ring series (*C. elegans-*specific edges are red, *P. pacificus*-specific edges are blue, core edges are black, and variable edges are gray). Legend applies to panels D-G. **E:** Histogram of directed connectivity conservation (Number of EM series with edge present) among all nematode nerve ring series. **F:** Histogram of adjacency conservation among adult nerve ring series. **G:** Histogram of directed connectivity conservation among adult nerve ring series. **H.** Violin plot comparing mean scaled adjacency and connectivity specificity classification Condition: No Connection (no statistical test performed), Not specific, U statistic: 541864, Corrected p-value: 3.11e-27, Condition: Core, U statistic: 3694, Corrected p-value: 1.80e-9 Condition: *P.* pacificus-specific, U statistic:2871, Corrected p-value: 7.56e-5 Condition: *C. elegans*-specific, U statistic: 180.0, Corrected p-value: 1.73e-4. Legend applies to panels H and I. **I.** Violin plot comparing log+1 adjusted number of synapses and connectivity specificity classification. Condition: not specific, U statistic: 1842058, Corrected p-value: 9.87e-67 Condition: core, U statistic: 1804.0, Corrected p-value: 2.72e-2 Condition: *P. pacificus*-specific, U statistic: 0.0, Corrected p-value: 2.33e-36 Condition: *C. elegans*-specific, U statistic: 12100.0, Corrected p-value: 1.02e-42. For underlying data, see **Supplement Data 5 and 6**.

If neuronal neighborhood shifts are uncommon, how widespread are small species-specific neighborhood rearrangements and do they instruct species-specific connections? Several above-mentioned extreme examples of changes in morphology (e.g. ASH, PVD or RIP) produce species-specific adjacencies associated with species-specific connectivity. In total, there are 28 instances of species-specific synaptic connections due to adjacencies that are exclusive to one specific species (**Figure 4 Supplement 1A, B).**

We undertook a more quantitative, whole-brain analysis of the relationship between neurite adjacency and synaptic connectivity. We assigned each neuron pair (edge) adjacency and synaptic conservation scores based on how frequently their connection appears across samples, with the score equal to the number of EM series containing that edge. We found that a greater proportion of adjacency edges are found in all *C. elegans* and *P. pacificus* samples than synaptic connectivity edges (**Figure 4D, E, Figure 4 Supplement 1B**). The same trend is found by comparing the adjacency (**Figure 4F**) and directed connectivity of only adult samples (**Figure 4G, Figure 4 Supplement 1A, B**). To contextualize these results, we compared the patterns conservation and specificity of adjacency and connectivity patterns across datasets to randomized null distributions. We found that our observed distributions of shared connections across datasets were higher than chance (**Figure 4 Supplement 1A**), and that the count of species-specific connections was statistically different from chance. This trend also holds for adjacencies, where we observed a greater number of shared adjacencies when compared to chance, with a statistically greater number of species-specific adjacencies observed. Together, these results show that the patterns of shared and species-specific adjacency and connectivity are much greater than chance, amplifying the importance of comparative analyses.

A previous analysis of neurite adjacency and synaptic connectivity revealed that the relative amount of neurite membrane adjacency predicts synaptic connections between neuron pairs in *C. elegans* (Cook et al. 2023). Consistent with this finding, we found that neuron pairs with species-specific connections are on average more strongly adjacent in that species than in the opposite species. (**Figure 4G**). For core connections, i.e., neuron pair connections that are conserved between the species, a smaller inter-species difference in mean scaled adjacency is observed (**Figure 4H**). This trend extends to synaptic connectivity, where we observed that the largest connections, on average, are ‘core’. Variable connections are on average, smaller than those found in one species (**Figure 4I**). Taken together, our analysis indicates that the core and species-specific connections are strongly related to the relative strength of neuronal adjacency.

Synaptic connectivity can be accurately modeled using relative adjacency in *C. elegans* (Cook et al. 2023). To explore whether the relationships between connectivity and adjacency are conserved, we extended this modeling approach to *P. pacificus.* We adjusted the training (*C. elegans* contact and connectomes) and test data (*P. pacificus* contact and connectomes), and used two predictors (mean scaled adjacency and brain strata)(Brittin *et al*. 2021) to classify synaptic connectivity (present in at least one sample). We found a moderately stronger relationship between mean scaled adjacency and mean connection weight among neuron pairs within vs. across different brain strata. (**Figure 4 Supplement 2A, B**) (Cook et al. 2023). Our modeling approach yielded superior results across classification algorithms (**Figure 4 Supp.1C**) compared to our previous results modeling connectivity in *C. elegans* alone. For comparative purposes, we plotted the Received Operator and Precision Recall curves for the Logistic Regression classifier, revealing an accurate connectivity prediction from simple neuronal adjacency characteristics (**Figure 4 Supplement 2D, E**). The striking similarity of our modeling results across species supports our previous proposal that relative adjacency is a guiding principle of synaptic organization in nematodes (Cook et al. 2023).

## Network configurations of the *C. elegans* and *P. pacificus* connectomes

Despite the importance of identifying similarities and differences in the parts list of the nervous system, we asked whether network topology differences exist between species. To first assess network consistency, we compared the degree (number of edges attached to a node) distributions of the *C. elegans* and *P. pacificus* chemical synaptic and adjacency networks. The degree distributions of the adjacency networks are strongly similar across datasets (**Figure 4 Supplement 4A)** with no statistically significant pairwise differences (**Figure 4 Supplement 4B).** Conversely, the synaptic degree distributions displayed widespread differences across development and species (**Figure 4 Supplement 4C, D)**. These findings support the notion that while the overall contactome structure remains stable, the connectome exhibits greater variability, being more influenced by the nuanced quantities of neuronal adjacency rather than by a binary presence-or-absence effect.

To better describe differences in network architecture across species we analyzed many commonly-used graph theoretical metrics across all datasets **(Figure 4 Supplement 5A).** There are many key organizational principles conserved across all datasets, including small-worldness, high clustering coefficients, and relatively short processing path lengths. We next performed hierarchical clustering using these graph metrics, revealing that networks cluster primarily by developmental stage **(Figure 4 Supplement 5B**). While transitivity and average clustering coefficients remained relatively stable across datasets, indicating the conservation of local processing modules, reciprocity values showed more variation, particularly in *P. pacificus,* suggesting species-specific differences in bilateral connectivity patterns. The degree assortativity coefficients were consistently negative across all networks, indicating that both species maintain a hierarchical hub-based organization where high-degree nodes tend to connect with low-degree nodes. Together, these results demonstrate that despite substantial evolutionary distance and different ecological niches, *C. elegans* and *P. pacificus* maintain similar fundamental organizational principles in their nervous systems, although *P. pacificus* may resemble mid-larval *C. elegans* in several ways.

To probe the putative functional units of the nervous system across species, we compared the motifs found in each connectomic dataset. We found strong differences between species where the motifs overrepresented in *C. elegans* were not overrepresented in *P. pacificus* **(Figure 4 Supplement 6A).** Two types of motifs, single-edge, and transitive triangles, were enriched in *P. pacificus* but not in *C. elegans.* The enrichment of transitive triangles (A→B→C, A→C) particularly indicates a prevalence of feed-forward processing with consistent hierarchical relationships between neurons. We speculate this architectural pattern might reflect *P. pacificus’s* more specialized behavioral repertoire, where direct, reliable signal propagation is prioritized. These findings suggest that while basic organizational principles are conserved between species, the relative emphasis on different circuit motifs has diverged through evolution to support species-specific behavioral requirements.

If specific types of circuit building blocks are different across species, could individual neurons be differentially important across species? To identify highly connected neurons that form the core of each connectome, we performed a conservative rich club analysis **(Figure 4 Supplement 7**). Rich clubs were identified using stringent criteria requiring both strong statistical enrichment (≥3 standard deviations above random networks) and robust topological features (≥3 consecutive degree thresholds). Several neurons with well-defined functional roles were identified as rich club members in both species: AIB, AVE, CEP, RIA, and RMD. RIP, AWA, AWB, and SAA were rarely or never rich club members in *C. elegans*, suggesting their differential centrality to the *P. pacificus* connectome. These graph theoretic findings, taken together through a conservative analytical approach, provide strong evidence for both conserved and species-specific aspects of core circuit organization in nematode nervous systems. Fundamental information processing differences have been maintained through evolution while allowing for species-specific adaptations in circuit architecture just as we observed in neuronal morphology.

## Glia connectivity as a substrate of evolutionary change

We also analyzed the other main component of the nervous system, glial cells. The ultrastructure of *P. pacificus* head glia supports a one-to-one correspondence of the three major cell types (sheath, socket, and GLR) (**Figure 5A, C, E**). Despite the anatomical similarity, we discovered one novel feature of *P. pacificus* GLR glia. These microglia-like cells extend two types of processes in both species: thin, leaf-like posterior processes that wrap the interior of the muscle-neuron plates of the nerve ring as well as anterior processes that fasciculate with inner labial dendritic sensory fascicles. (**Figure 5C,F**)(Hong *et al*. 2019). Strikingly, in *P. pacificus,* these anterior extensions contain bona fide presynaptic features, e.g., swellings, active zones and synaptic vesicles (**Figure 5G**). The GLR cells receive synaptic input from RIP in both species but make *P. pacificus*-specific output onto inner- and outer labial cells, other glia and the pharyngeal epithelium (**Figure 5H**). So-called “gliotransmission” has been reported in vertebrate astrocytes (Savtchouk and Volterra 2018) but has not yet been revealed as a substrate of evolutionary change in a nervous system. As we already speculated in the context of the intriguing wiring differences of the RIP neurons, the ability of *P. pacificus* GLR glia to communicate with pharyngeal tissue may relate to predatory feeding behavior.

**Figure 5:**
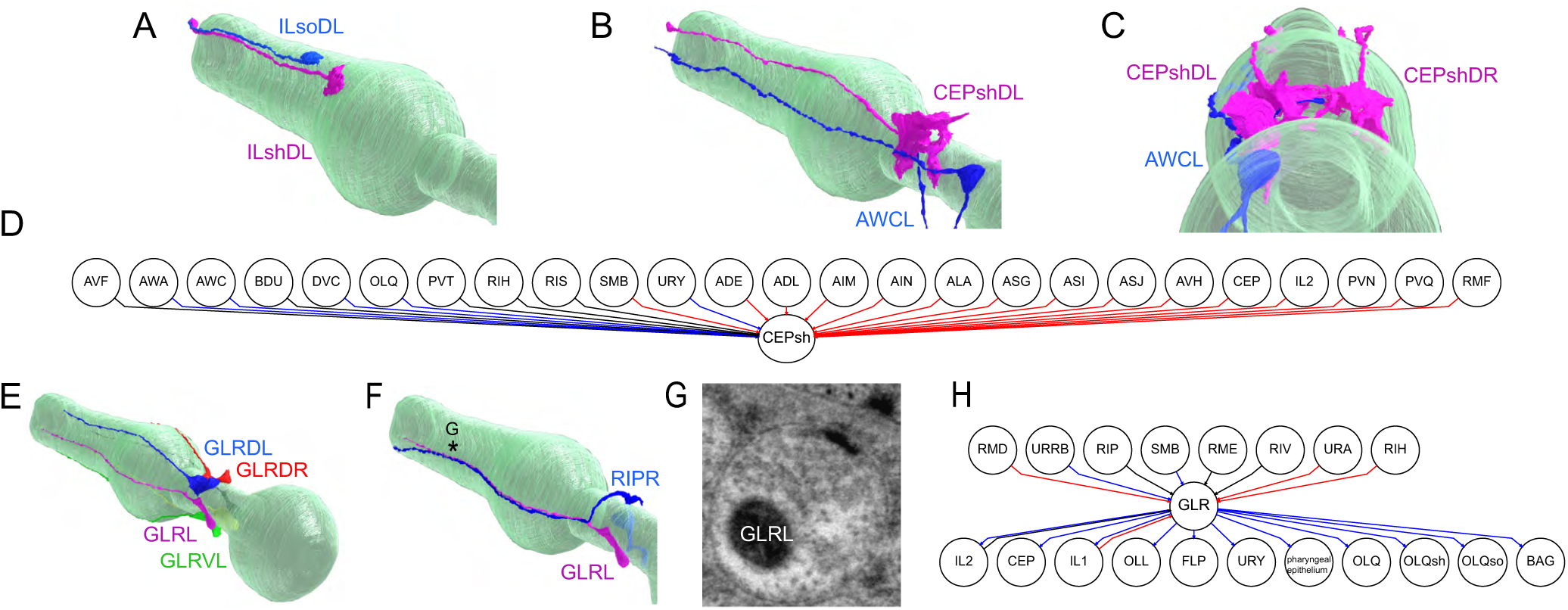
Glial rewiring across species. **A:** Lateral posterior view of 3D rendering of IL sheath and socket cell **B:** Lateral posterior view of 3D rendering of the CEPshDL and AWCL cells. **C:** Posterior view rendering of CEPshDL/R and AWC cells. **D:** Chemical synaptic connectivity of CEPsh in adults. Blue lines are connections found only in *P. pacificus*, red lines are connections found only in *C. elegans*, and black lines are connections found in both species. **E:** Lateral posterior view rendering of 3D reconstruction of the GLR cells. **F:** Posterior lateral view of the GLRL and RIPL cells, showing close fasciculation of processes. **G:** Region of synaptic output of GLR cells (pink) with an EM micrograph showing presynaptic specializations within the GLR cell **H:** Chemical synaptic connectivity of GLR in adults. Blue lines are connections found only in *P. pacificus*, red lines are connections found only in *C. elegans*, and black lines are connections found in both species.

We discovered that the astrocyte-like CEPsh glia also displays striking changes in synaptic connectivity. The extensions of the CEPsh glia that line the nerve ring are innervated by the neurites of several distinct neuron types in the *C. elegans* nerve ring (Witvliet *et al*. 2021). Like the GLR glia, the *P. pacificus* CEPsh glia are localized to nearly identical anatomical locations with no gross shift in neighborhood placement. In *P. pacificus*, several of these inputs are lost and others are gained. For example, there is now strong innervation of the CEPsh glia by the AWC olfactory neurons in the nerve ring (**Figure 5B, C, D**). Vertebrate astrocytes also receive synaptic inputs that are translated into gliotransmitter release (Savtchouk and Volterra 2018). While we observe no obvious synaptic release sites in the CEPsh glia, we note that at least in *C. elegans*, the CEPsh glia release neuropeptides to control stress resistance and animal lifespan (Frakes *et al*. 2020). If this CEPsh function is preserved in *P. pacificus*, a rewiring of CEPsh glia indicates that the neuronal control of such important physiological parameters may be subject to evolutionary change.

## Conclusions

The exceptionally well-described neuronal anatomy of *C. elegans* combined with our analysis of multiple *P. pacificus* datasets yields a comprehensive map of how brains change over millions of years of evolution. Our analysis revealed multi-tiered substrates of evolutionary change, ranging from alteration in the number of individual constituent neurons (exemplified by species-specific neuronal cell death), cell body position, projection patterns of dendrites and/or axons, to changes in synaptic connectivity which result in novel network structures.

Brain development in vertebrate and invertebrate animals, including nematodes, involves a substantial amount of cell death (Hengartner and Horvitz 1994; Fricker *et al*. 2018). The discovery of alterations in the cellular and temporal specificity of neuronal cell death programs in *C. elegans* and *P. pacificus* indicates that programmed cell death may constitute an important driving force in shaping circuit architecture over evolutionary timescales. Such a mechanism may explain the observation that in several nematode species more or fewer inner or outer labial sensory neurons, have been observed (Endo 1980; Fine *et al*. 1997; Ragsdale *et al*. 2009); in *C. elegans* sisters of these labial sensory neurons undergo cell death (Sulston *et al*. 1983) and, hence, a precocious or a lack of execution of the cell death program in these lineages compared to *C. elegans* could explain these differences in cellular composition of the labial sensory apparatus. Changes in neuronal composition through altered cell death patterns have been observed in related *Drosophila* species (Pop *et al*. 2020; Prieto-Godino *et al*. 2020), indicating that this mechanism of brain evolution is conserved across phylogeny.

Apart from providing the most comprehensive definition of a genus’s conserved connectome to date, we found that most neurons generate species-specific synaptic connectivity, arguing for widespread changes in information flow and processing. Intriguingly, changes in network structure do not necessarily result in behavioral changes, as illustrated by the ASH sensory neurons which, based on their synaptic connectivity, appear to transmit sensory information to a distinct set of downstream interneurons. Yet, in both species, ASH triggers nociceptive responses to aversive cues (Kaplan and Horvitz 1993; Srinivasan *et al*. 2008). Akin to the concept of developmental systems drift (True and Haag 2001), the ASH circuit may constitute an example of “circuit drift”, in which the pathways of information processing may have drifted, while keeping behavioral output constant (Katz 2016; Roberts *et al*. 2022).

Changes in synaptic connectivity are largely, but not exclusively, the result of changes in relative process adjacency, which supports the applicability of Peters’ Rule for the generation of synaptic specificity in nematodes (Cook *et al*. 2023). We speculate that at least some of the changes in circuit architecture relate, in particular, the rewiring of RIP, to the intriguing predatory behavior that *P. pacificus* can engage in.

Apart from revealing the specific substrates of evolutionary change, the analysis of whole brains (rather than isolated parts) permitted us to address the fundamental question of whether evolutionary changes homogenously distribute across the network or are concentrated at specific hotspots, such as the sensory periphery. Our analysis demonstrates that the evolutionary conservation of synaptic connectivity and evolutionary novelty are indeed distributed throughout the entire brain.

## METHODS

### EM analysis methodology

Two serial section EM series of eurystomatous (predatory) hermaphrodite animals (Bumbarger *et al*. 2013) from the PS312 strain were further reconstructed for ultrastructural morphology and connectivity using TrakEM2 (Cardona *et al*. 2012). In brief, each neuron profile present in an EM section was manually traced to create volumetric neuron reconstructions. From these series, ROIs were created which correspond to the nerve ring and surrounding neuropil. We limited our comparative analysis of neuronal and adjacency to this ROI, but report the connectivity data in **Supplement Data 1,2**. Neural adjacency data for *P. pacificus* were computationally extracted as previously described (Brittin *et al*. 2021), with slight modification to extract ‘areatree’ annotations using multiprocessing. We manually identified chemical synapses and gap junctions between neurons and tissues using criteria established previously (White *et al*. 1986; Cook *et al*. 2019), but excluded gap junction connectivity from our comparative analysis due to ultrastructural ambiguity, a persistent issue in samples processed for connectomics. Connectivity data were extracted from TrakEM2 using a combination of built-in tools and custom python parsing scripts. These extracted connectivity data include the number of synapses and weight metric (number of consecutive serial sections where synaptic specializations are observed) for each EM series. Our analysis is limited to the number of connections between two neurons unless noted otherwise.

### Randomization and graph theoretic analysis

Synaptic connectivity and adjacency data were compiled across all datasets such that each row represents a neuron pair and its presence or properties are stored in columns (e.g. number of shared datasets, strata membership, average synaptic size). To generate randomized null distributions, we randomly swapped edges between neuron pairs preserving the overall degree distribution for 5x the number of edges in each network, repeating for 10,000 iterations. At each iteration, we calculated the distribution of edge counts and species-specific edges found. We then compared these distributions to the observed data, calculating a Z-score and two-tailed p-value for each.

### Assignment of homologous neurons

We limited our cross-species comparison of ultrastructure to normal *C. elegans* development and only the hermaphrodite sex. Given its rich description of inter-individual variability, we used anatomical landmarks of *C. elegans* as a reference. To identify neuronal identities, we classified cells based on the following features (in order of importance): relative soma location, symmetry (two-fold, four-fold, six-fold), soma size, number and direction of processes, hallmark structural features (e.g. ‘humped’ cross-sectional morphology of AIY in the ventral ganglion) as previously described (White *et al*. 1986; Cook *et al*. 2019). Further ambiguities were resolved by evaluating the specific location of a process within the neuropil and distinct swellings in axons or dendrites. In extreme cases, such as the AVH neuron, embryonic lineaging resolved any ambiguity. We avoided the use of synaptic connectivity to identify cells. All neuronal cells with connected somas were identified within our region of interest (ROI). However, because the EM series did not cover the entire animal, we were unable to positively identify all processes within the ROI and, therefore, excluded those from our analysis. These cells include ‘unknown’ RVG neurons that project into the ventral ganglion, which do not have homologous counterparts in *C. elegan*s. Additionally, some axons project from the posterior of the animal into the nerve ring but lack distinguishing characteristics in *P. pacificus* which are seen in *C. elegans*, such as prominent neighborhood positions or distinct branching patterns. For example, there is a bilateral pair of *P. pacificus* axons that fasciculate in the nerve ring similar to HSN in *C. elegans*, but unlike *C. elegans* these *P. pacificus* neurons are unbranched and both travel in the right fascicle of the ventral nerve cord. Serotonin staining patterns also suggest no obvious HSN-like neuron in the adult *P. pacificus* (Loer and Rivard 2007).

The pharyngeal and amphidial circuits were previously homologized with a 1:1 correspondence between each species in their neuronal identity (Bumbarger *et al*. 2013; Hong *et al*. 2019). Due to details of their axonal projection patterns, the analysis of reporter data and discussions with R. Hong, we exchange the identities of *P. pacificus* ASE and AWC in this paper relative to a previous analysis of the amphid circuitry of *P. pacificus* (Hong *et al*. 2019). To make accurate morphological comparisons, we evaluated the morphology of P. pacificus neurons in comparison to all available 3D reconstructions in C. elegans (White *et al*. 1986; Brittin *et al*. 2021; Witvliet *et al*. 2021).

### Machine learning assumptions, selection, performance

Our methodology for assumptions, selection, and performance of classification algorithms was used as previously described (Cook *et al*. 2023). In brief, we classify each edge (pairs of neurons known to be adjacent) as a binary value for synaptic weight (>0 average connection size = 1, no connection observed across samples = 0). We then trained our model using *C. elegans* adjacency and connectivity data using the mean scaled and normalized adjacency of pairs and a categorical variable describing whether two neurons are in the same or different strata (Brittin *et al*. 2021). Our models (MLPClassifier, DecisionTreeClassifier, RandomForestClassifier, and LogisticRegression from the scikit-learn library (Pedregosa *et al*. 2011) were then evaluated on the test dataset of *P. pacificus* adjacency and connectivity data. For comparative purposes, we again chose the parameters previously used in our logistic regression model for evaluating *C. elegans* adjacency and connectivity data.

### Visualizing morphology and circuitry

Renderings of individual neurons or groups of neurons from Specimen 104 were performed using Adobe Dimension or Blender. Some post-rendering modifications of images were done with GraphicConverter. All code required to automate the processing of Blender data are available at https://github.com/stevenjcook/cook_et_al_2024_mipristionchus. Individual neuron renderings, Blender and Adobe Dimension files used for rendering will be made available on Zenodo upon publication.

Circuit diagrams for individual neuron classes were generated using GraphViz (https://graphviz.org/) using the ‘circo’ layout. Display layouts of complete networks were made using Cytoscape (Smoot *et al*. 2011). The layout structure is an implementation of an affinity view popularized by (Varshney *et al*. 2011).

### Quantification and statistical analysis

The statistical analysis performed in this paper is a combination of python software packages which include Scikit-learn and SciPy. All information for individual statistical tests including test and p-values can be found in the figure legends, while summary statistics are reported in the main test. All graph theoretic analysis code is available on Github with reproducible figures.

### 4D-microscopy and lineage analysis

The method for 4D-microscopy was described previously (Schnabel et al. 1997). Modifications of this system are described in (Schnabel *et al*. 2006). All 4D recordings generated were analyzed using the Software Database SIMI©BioCell (SIMI Reality Motion Systems, Unterschleissheim, Germany; http://www.simi.com/)(Schnabel *et al*. 1997; Schnabel *et al*. 2006). Cells are followed by the observer and the coordinates are recorded approximately every 2 min. The cell cleavages are assessed by marking the mother cell before the cleavage furrow ingresses and subsequently the centers of the daughter cells three frames later. By marking every cell during the complete embryonic development, the complete cell lineage of an embryo is generated.

### Resource availability

This study did not generate new or unique reagents. Additional requests for information related to data analysis should be directed to SJC and OH.

### Data Sources

*C. elegans* nerve ring connectivity and adjacency data were previously published (Cook *et al*. 2019; Brittin *et al*. 2021; Witvliet *et al*. 2021). The *P. pacificus* electron micrographs used in this study were previously published (Bumbarger *et al*. 2013) and previously analyzed for amphidial circuitry (Hong *et al*. 2019).

## Supporting information

Supplementary Figues

## ACKNOWLEDGEMENTS

This work was funded by the Howard Hughes Medical Institute. We thank Y. Ramadan, N. Schroeder, R. Hong, and M. Chalfie for discussions of *P. pacificus* gene expression, behavior and anatomy and David Hall, James Lightfoot, Nathan Schroeder, and members of the Hobert lab for comments on the manuscript.

## AUTHOR CONTRIBUTIONS

SJC – conceptualization, methodology, investigation, formal analysis, software, writing, editing

CAK – investigation, formal analysis, software

CML – investigation, formal analysis, visualization, editing

NM – investigation, formal analysis

MM – investigation

SRS – investigation

DJB – conceptualization, methodology, investigation

MR – methodology, investigation

RS – investigation, formal analysis

RJS – editing, supervision, funding acquisition

OH – conceptualization, investigation, writing, editing, funding acquisition, supervision

## DATA AVAILABILITY

All neuronal adjacency and connectivity data necessary for the analyses presented in this paper are included in Supplement Data 1-6.

## CODE AVAILABILITY

All code necessary to analyze neuronal adjacency and connectivity data in this paper are provided are available at https://github.com/stevenjcook/cook_et_al_2024_pristionchus.

