## Supplementary Figues for "Comparative connectomics of two distantly related nematode species reveals patterns of nervous system evolution"

### SUPPLEMENTAL DATA, FILES AND FIGURES

**Supplement Data 1:** Neuronal adjacencies within the nerve ring of Specimen 107.
**Supplement Data 2:** Neuronal adjacencies within the nerve ring of Specimen 148.
**Supplement Data 3**: Connectivity of *P. pacificus* reconstruction from Specimen 107
**Supplement Data 4**: Connectivity of *P. pacificus* reconstruction from Specimen 148.
**Supplement Data 5:** Composite dataset of undirected adjacency and connectivity across nematode samples
**Supplement Data 6:** Composite dataset of directed adult chemical connectivity across nematode samples.

**Supplement File 1: 3D renderings of all reconstructed individual neurons.**
**A:** Somatic neurons. **B:** Pharyngeal enteric neurons.
3D renderings from 'Series 14' of all reconstructed individual *P. pacificus* neurons, each named for the presumptive *C. elegans* homologous neuron. Most neurons are shown with their bilateral homolog; the left cell in magenta, right blue. This includes 4 and 6-fold symmetric neurons. In one case, a dorsal – ventral pair is shown (RMED, RMEV). Unpaired neurons are shown alone. All head neurons are shown rendered in multiple views (see below). As a constant landmark, the pharynx is shown in light green. Disrupted neuron trajectories in the posterior reflect that the serial section electron micrograph series ends within the retrovesicular ganglion. In all images, anterior is to the left. Viewpoints, top to bottom; views 1 – 4 views are rotated ~90° each step approximately around the pharyngeal center on the anterior-posterior axis: 1. Left lateral (top), 2. Dorsal, 3. Right lateral, 4. Ventral, 5. Rear, left side (bottom), anterior to the upper left. For both left (1) and right (3) lateral views, the view is slightly tilted relative to being precisely lateral, to better show the opposite bilateral homolog for paired neurons. The relative 'zoom' of different renderings varies to show the complete set of neurites and/or soma locations. The viewpoints are most 'zoomed out' for cells with somas and/or neurites in the retrovesicular ganglion. Rendered with Adobe Dimension from .obj files exported from TrakEM2. Some post-rendering retouching was performed with Graphic Converter, especially changing the background grey to enhance edge contrast.

**Supplement File 2: Directed connectivity of all head neurons shown individually.** For each neuron reconstructed in our datasets, a circuit diagram is presented where variable connections are shown in gray, core connections are shown in black, *C. elegans*-specific connections in red, and *P. pacificus*-specific in blue. One neuron is shown per page.

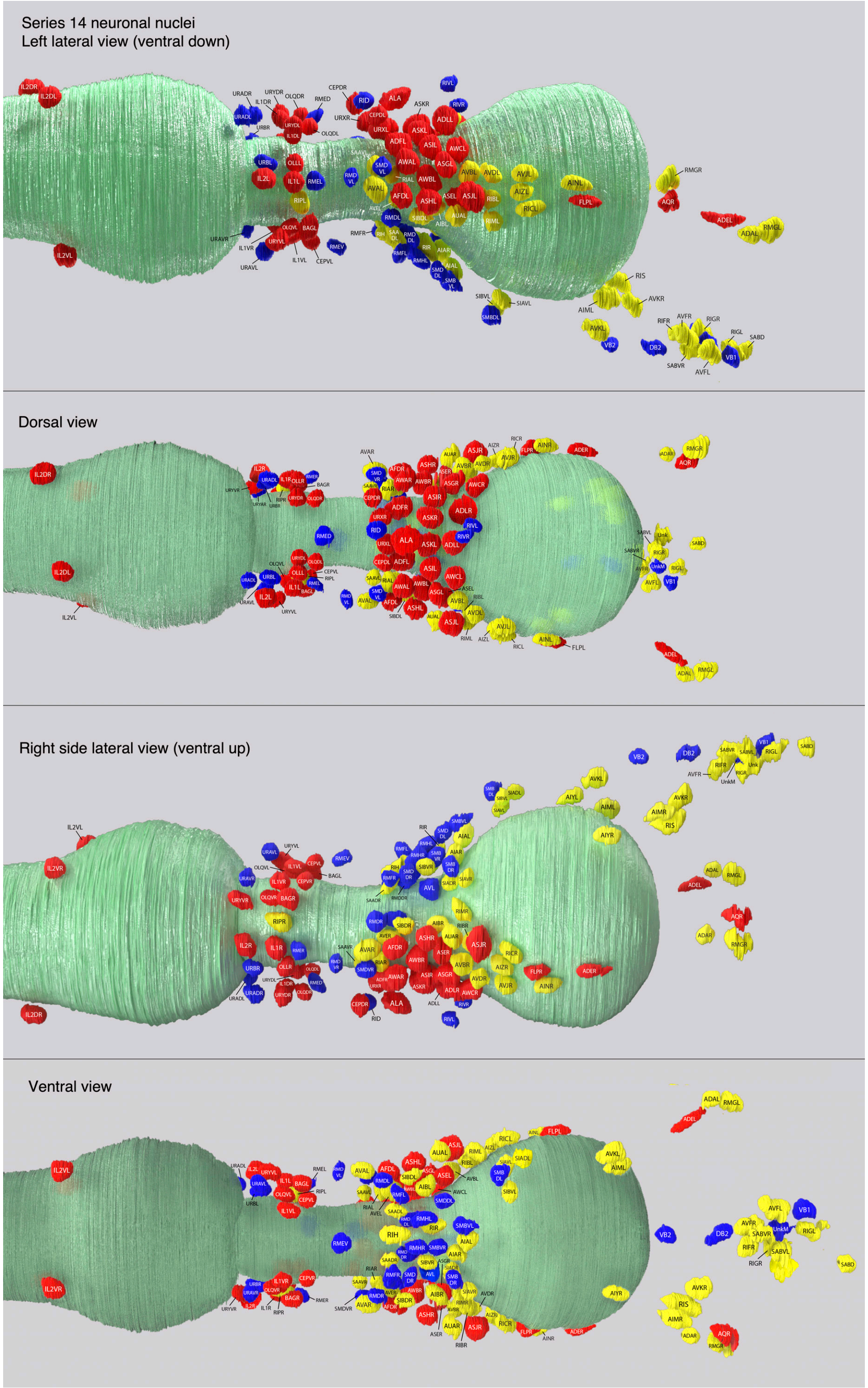


**Figure 1 - SUPPLEMENT 1: Neuronal cell body positions.** Different perspectives of neuronal cell body position.

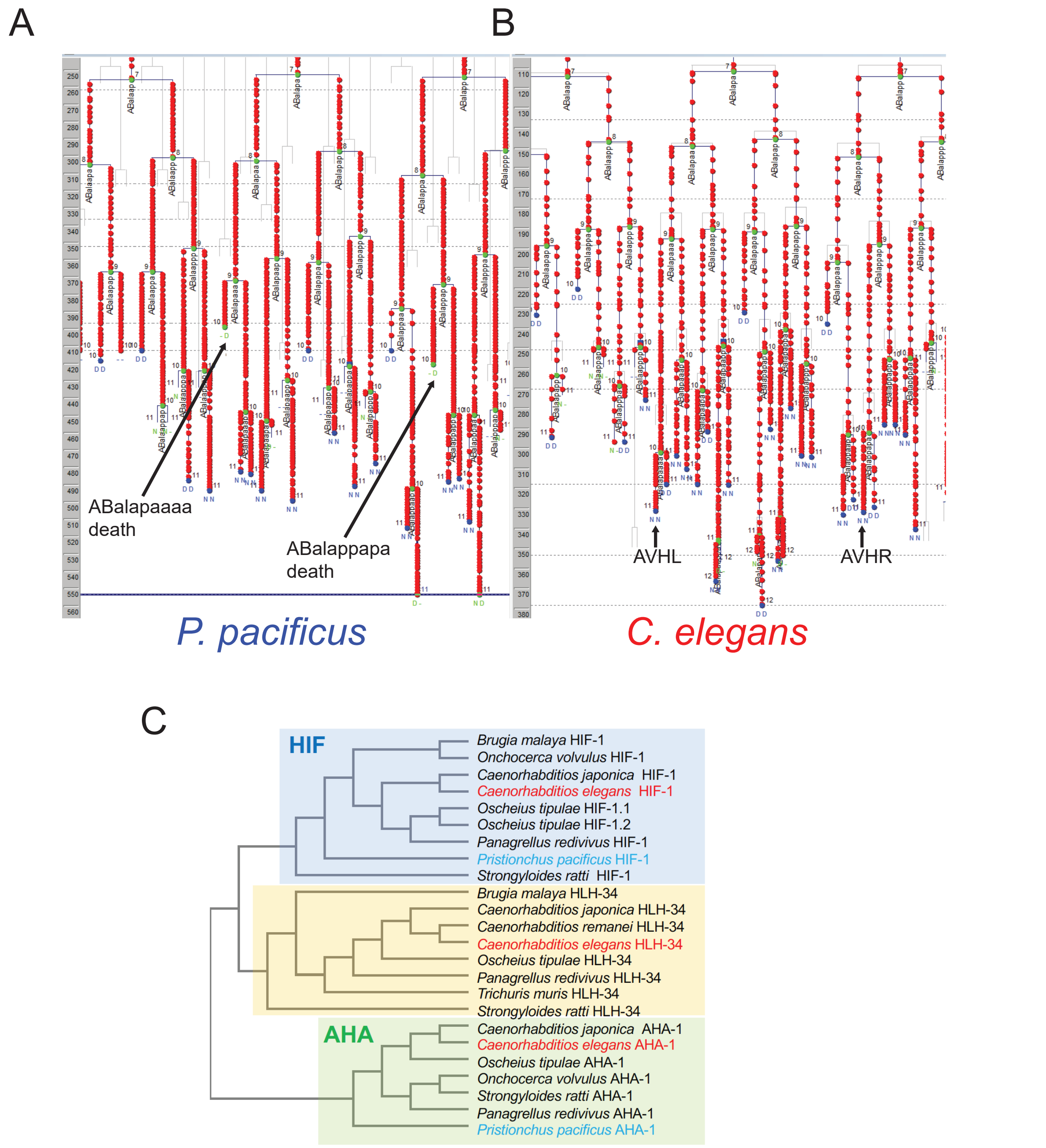


**Figure 1 - SUPPLEMENT 2: AVH neuron's progenitor dies in *P. pacificus,* but not in *C. elegans.***

Primary lineage data generated by Simi BioCell is shown. Cell fates are defined by nuclear morphology (N= neuron-like nuclear morphology; D = cell death; U = muscle; M = mitosis; P = pharynx; H = hypodermal; duplicated letter means: first letter = predicted fate; second letter = observed fate).

**A:** Lineage data from the *P. pacificus* embryo, showing the death of AVHL/R's mother cell.

**B:** Lineage data from the *C. elegans* embryo, highlighting AVHL/R.

**C:** Phylogenetic tree of the PAS domain of bHLH-PAS proteins from various nematode species, illustrating the loss of *hlh-34* and highlighting *P. pacificus* (blue) vs *C. elegans* (red).

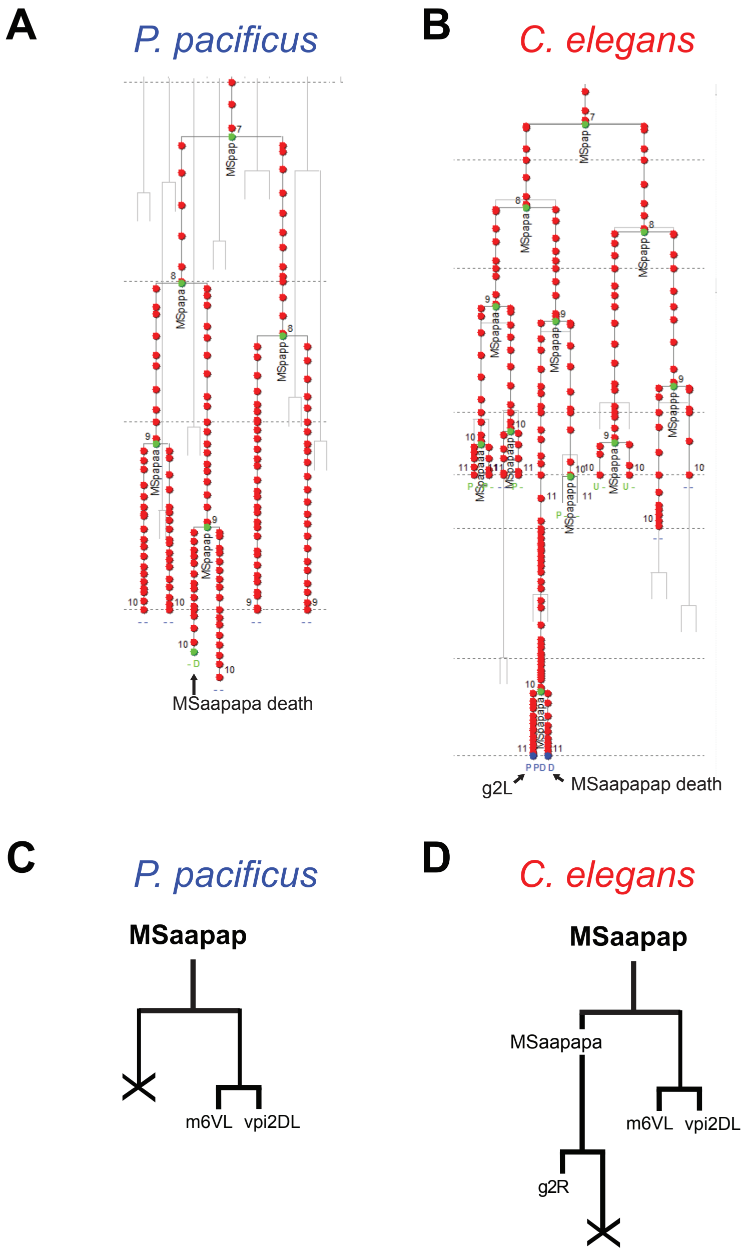


**Figure 1 - SUPPLEMENT 3: g2 gland progenitor dies in *P. pacificus,* but not in *C. elegans.***

Primary lineage data generated by Simi BioCell is shown. Cell fates are defined by nuclear morphology (See previous legend for the letter code).

**A:** Lineage data for the *P. pacificus* embryo, showing the death of g2L's mother cell.

**B:** Lineage data for *C. elegans* embryo, highlighting the death of g2L's sister cell.

**C:** Schematic lineage diagram depicting the death (cross) of g2L's mother cell in *P.*

*pacificus*.

**D:** Schematic lineage diagram depicting the death of g2L’s sister cell (cross) in *C. elegans.*

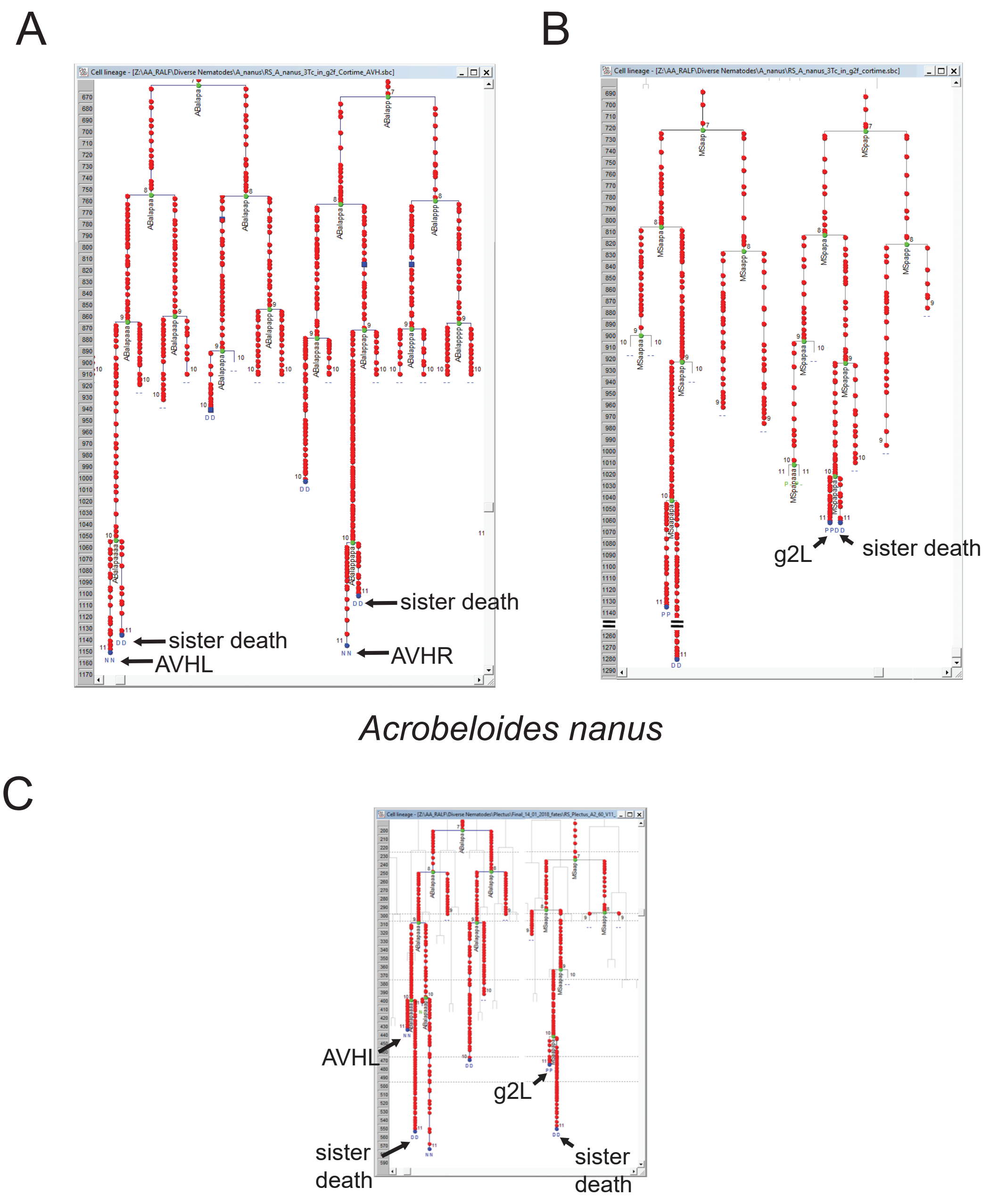


*Plectus sambesii*

**Figure 1 - SUPPLEMENT 4: AVHL and g2 survive in *Acrobeloides nanus* and *Plectus sambesii*, as in *C. elegans.***

Primary lineage data generated by Simi BioCell is shown. Cell fates are defined by nuclear morphology (See previous legend for the meaning of the letter code).

**A:** Lineage data for the *A. nanus* embryo, showing the death of AVHL's sister cell (left) and AVHR's sister cell (right).

**B:** Lineage data for the *A. nanus* embryo showing the death of g2L's sister cell.

**C:** Lineage data for the *P. sambesii* embryo showing the death of the sister cells of AVHL and g2L.

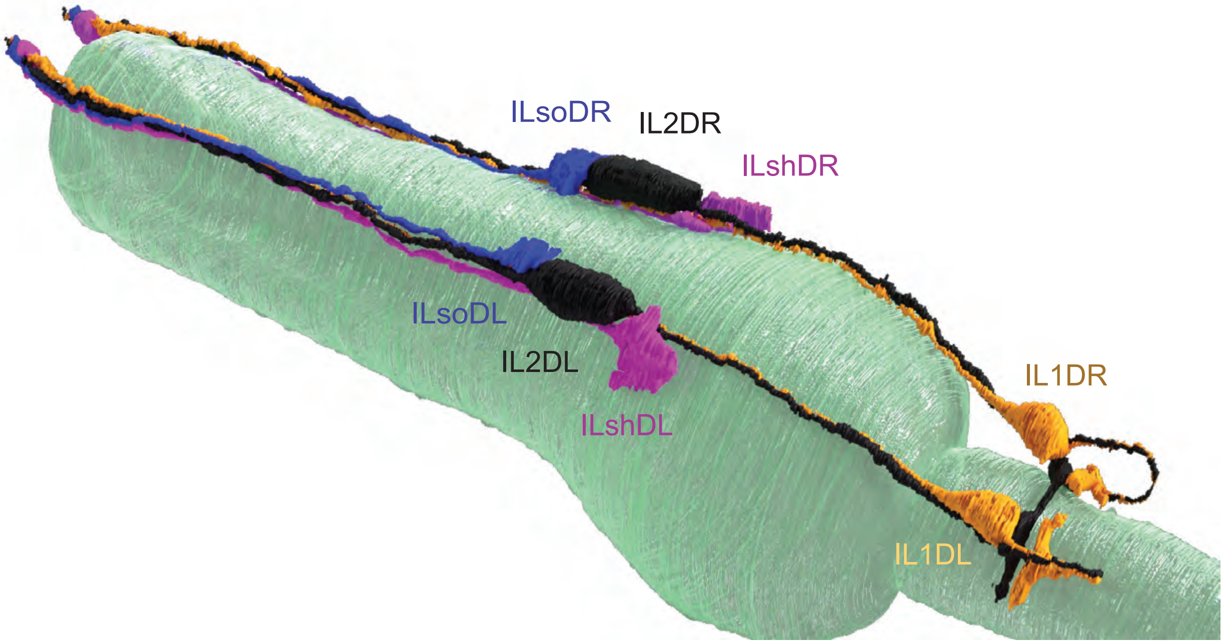


**Figure 1 - SUPPLEMENT 5: Anterior displacement of the IL2 neuron somas**. Rendering of the dorsal IL2 neuron pair (black). Its axons are closely fasciculated with the IL1 neuron pair (orange) processes. The IL2D somas are nestled between the IL socket (blue) and IL sheath (magenta) glia somas, with the pharynx shown in teal.

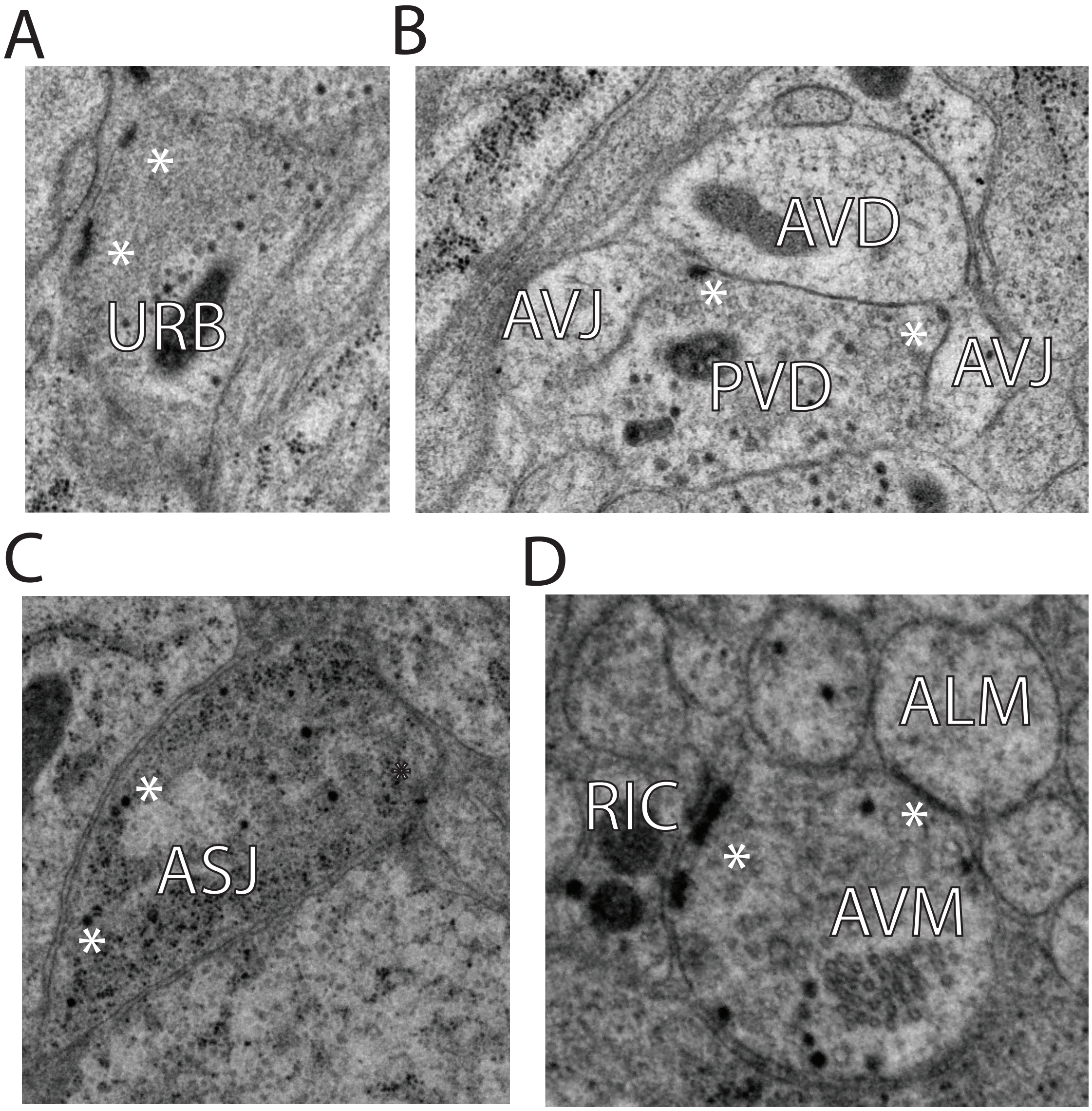


**Figure 2 - SUPPLEMENT 1: Ultrastructural details of anatomical differences.**

**A:** EM micrograph showing URB NMJs. Active zones are shown with asterisks.

**B:** EM micrograph showing PVD's synaptic output onto the AVJ and AVD neurons. Active zones are shown with asterisks.

**C:** EM micrograph showing dense core vesicles within ASJ's species-specific perisomatic branch.

**D:** EM micrograph showing a chemical synapse (left asterisk) and gap junction (right asterisk) along AVM's microtubule-containing process.

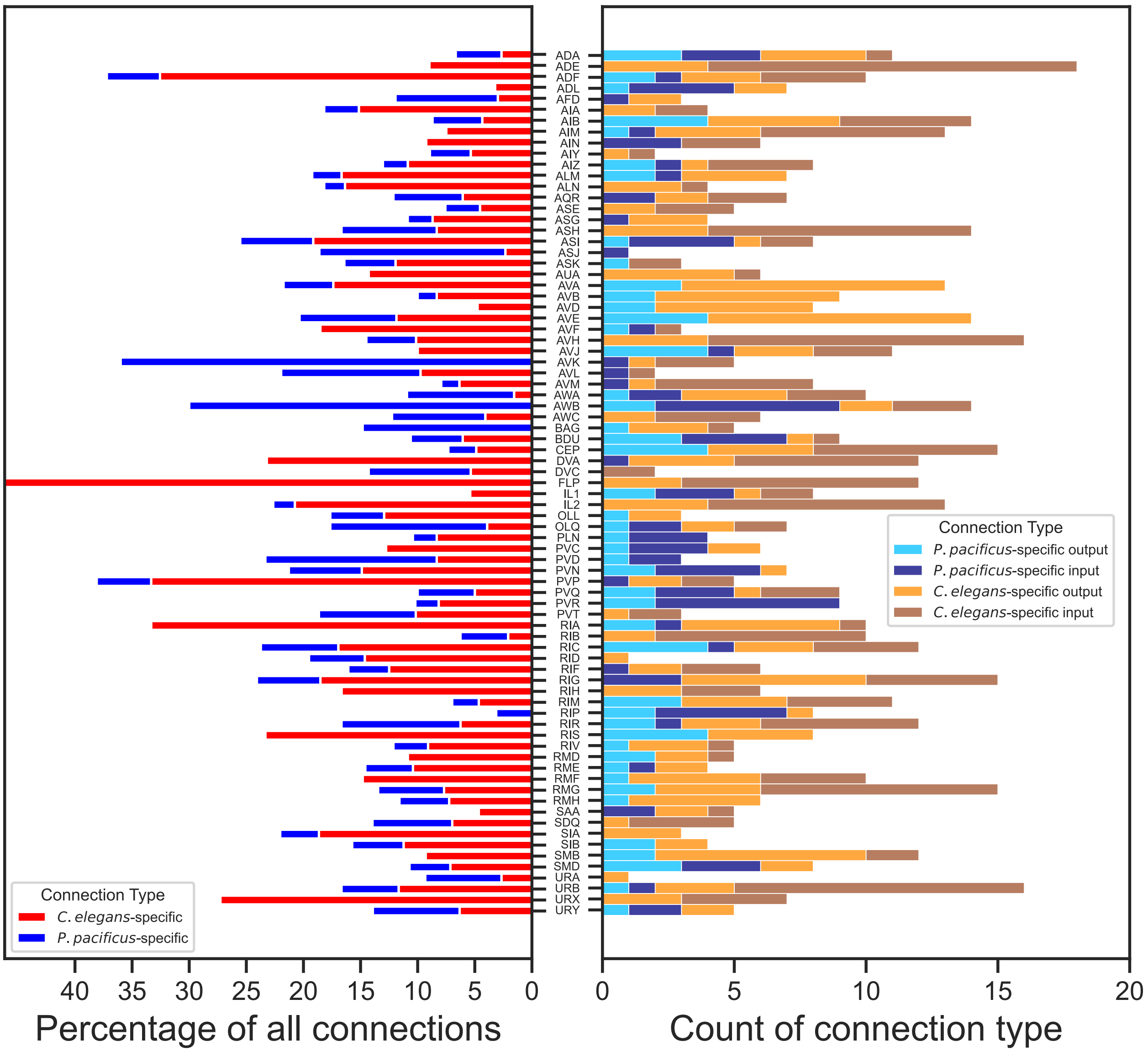


**Figure 3 – SUPPLEMENT 1: Analysis of species-specific adult chemical connectivity**

All embryonically born sex-shared nerve ring neurons that make a species-specific synaptic connection are shown. Stacked bar graphs of the percentage of species-specific undirected synaptic connections relative to all other synapses made by the respective neuron class (left).Stacked bar graphs of the number of sex-specific directed inputs and outputs by neuron class (right).

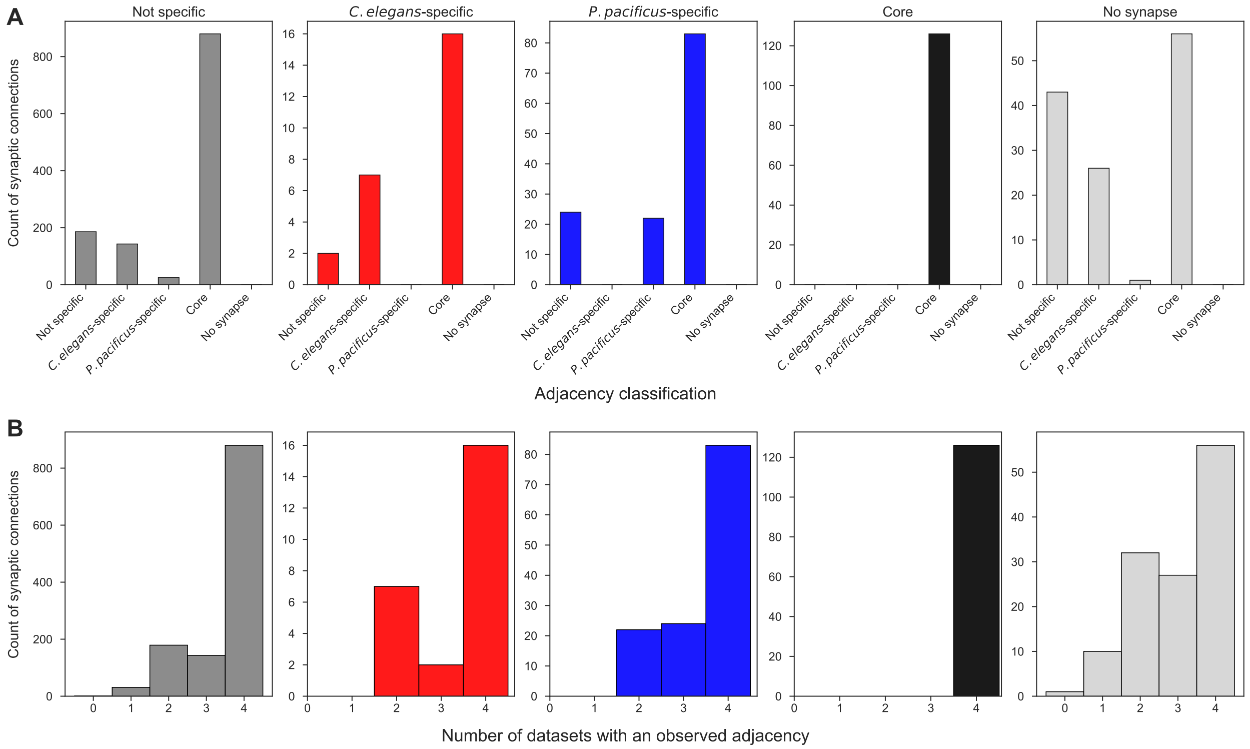


**Figure 4 - SUPPLEMENT 1: Characteristics of adult synaptic connectivity.**

**A:** Bar graphs of the adult synaptic connectivity classification (facets) vs adjacency classification (x-axis ticks).

**B:** Histograms of the count of synapses by connectivity classification (facets) compared to the number of adult EM series with an observed adjacency.

A

B

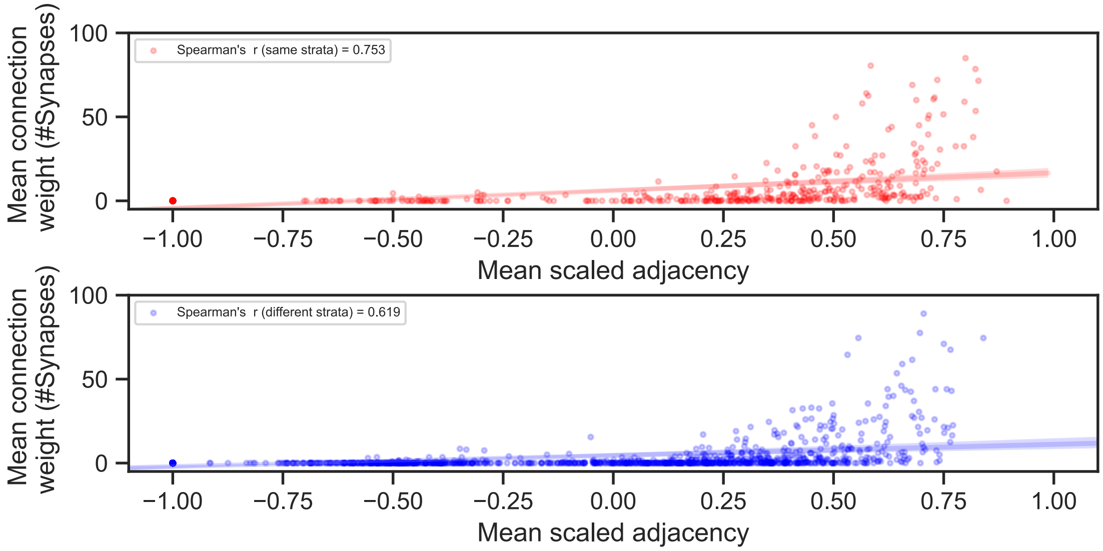


C

| Metric | Multi-layer Perceptron | Random Forest | Decision Tree | Logistic Regression |
|---|---|---|---|---|
| Area Under The ROC Curve | 0.8558 | 0.8535 | 0.8492 | 0.8511 |
| Area Under The Precision Recall Curve | 0.8577 | 0.837 | 0.8281 | 0.8508 |
| F1 Score | 0.7631 | 0.7723 | 0.732 | 0.7624 |

D

E

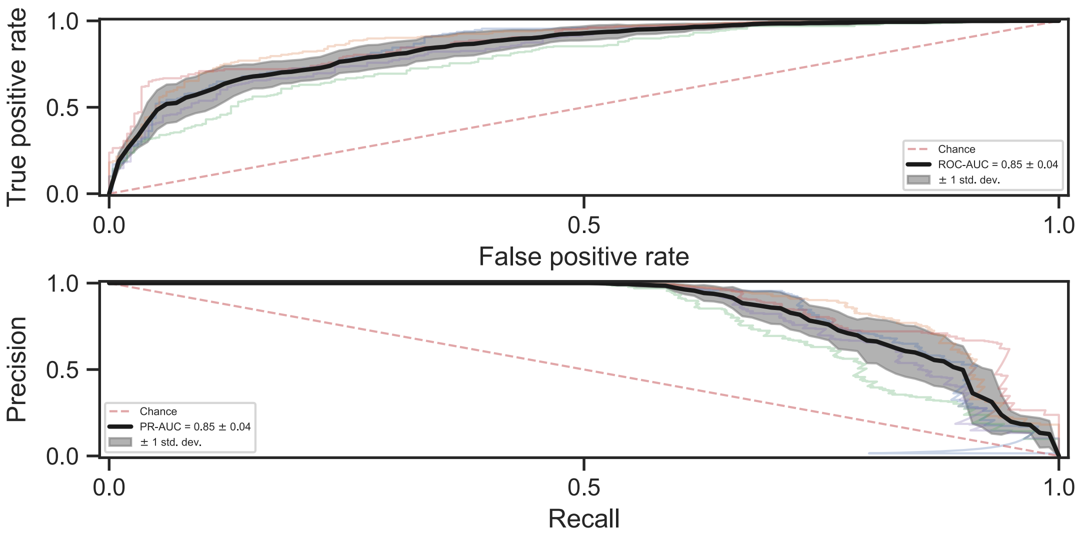


**Figure 4 - SUPPLEMENT 2: Neuronal adjacency predicts connectivity in *P. pacificus.***

**A.** Red scatter plot of the mean scaled adjacency vs. mean connection strength for neuron pairs in the same strata. The regression line is shown in red, Spearman's r =

0.7753

**B.** Blue scatter plot of mean scaled adjacency vs. Mean connection strength for neuron pairs in different strata. The regression line is shown in blue, Spearman's r = 0.619.

**C.** Table of 10-fold cross-validation of Multi-layer Perceptron, Random Forest, Decision Tree, and Logistic Regression classification algorithm results.

**D.** Receiver operator curves of the cross-validated model (kfolds = 10). The black line and gray area are the mean (0.85) and standard deviation (0.04), respectively.

**D.** Precision recall curves of the cross-validated model (kfolds = 10). The black line and gray area are the mean (0.85) and standard deviation (0.04), respectively.

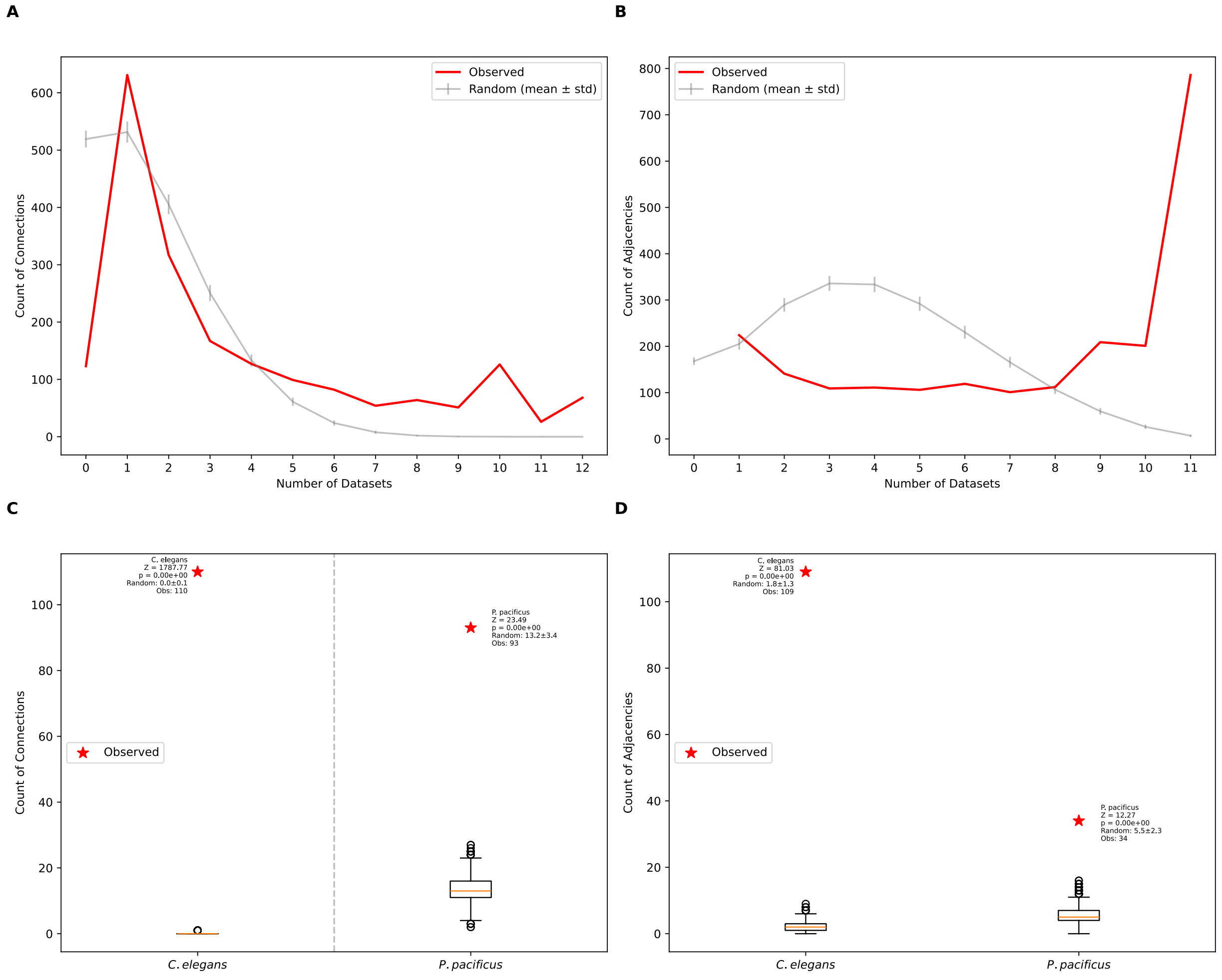


**Figure 4 SUPPLEMENT 3: Comparison of observed synaptic connectivity and adjacency patterns to randomized null distributions.**

**A**: Distribution of the number of directed synaptic connection edges shared across datasets compared (red) to randomized networks preserving degree distribution (gray, mean ± std).

**B:** Distribution of the number of undirected neuronal adjacency edges shared across datasets compared (red) to randomized networks preserving degree distribution (gray, mean ± std).

**C:** Number of synaptic connections specific to *C. elegans* (present in all *C. elegans* datasets but zero *P. pacificus* datasets) and specific to *P. pacificus* (present in both P. pacificus datasets but zero C. elegans datasets) compared to randomized distributions. *C. elegans* has significantly more species-specific connections (110) than expected by chance (Z = 1787.77, p = 0.00), as does *P. pacificus* (93, Z = 23.49, p = 0.00).

**D:** Number of neuronal adjacencies specific to *C. elegans* (present in all *C. elegans* datasets but zero *P. pacificus* datasets) and specific to *P. pacificus* (present in both P. pacificus datasets but zero C. elegans datasets) compared to randomized distributions. *C. elegans* has significantly more species-specific adjacencies (109) than expected by chance (Z = 81.03, p = 0.00), as does P. pacificus (34, Z = 12.27, p = 0.00). Z-scores were calculated as (observed value - random mean) / random std. Two-tailed p-values were calculated as 2 (1 - CDF of abs(Z)) for the standard normal distribution. n = 1000 randomized networks for each comparison. Boxplots show median, interquartile range, and 1.5 IQR whiskers.

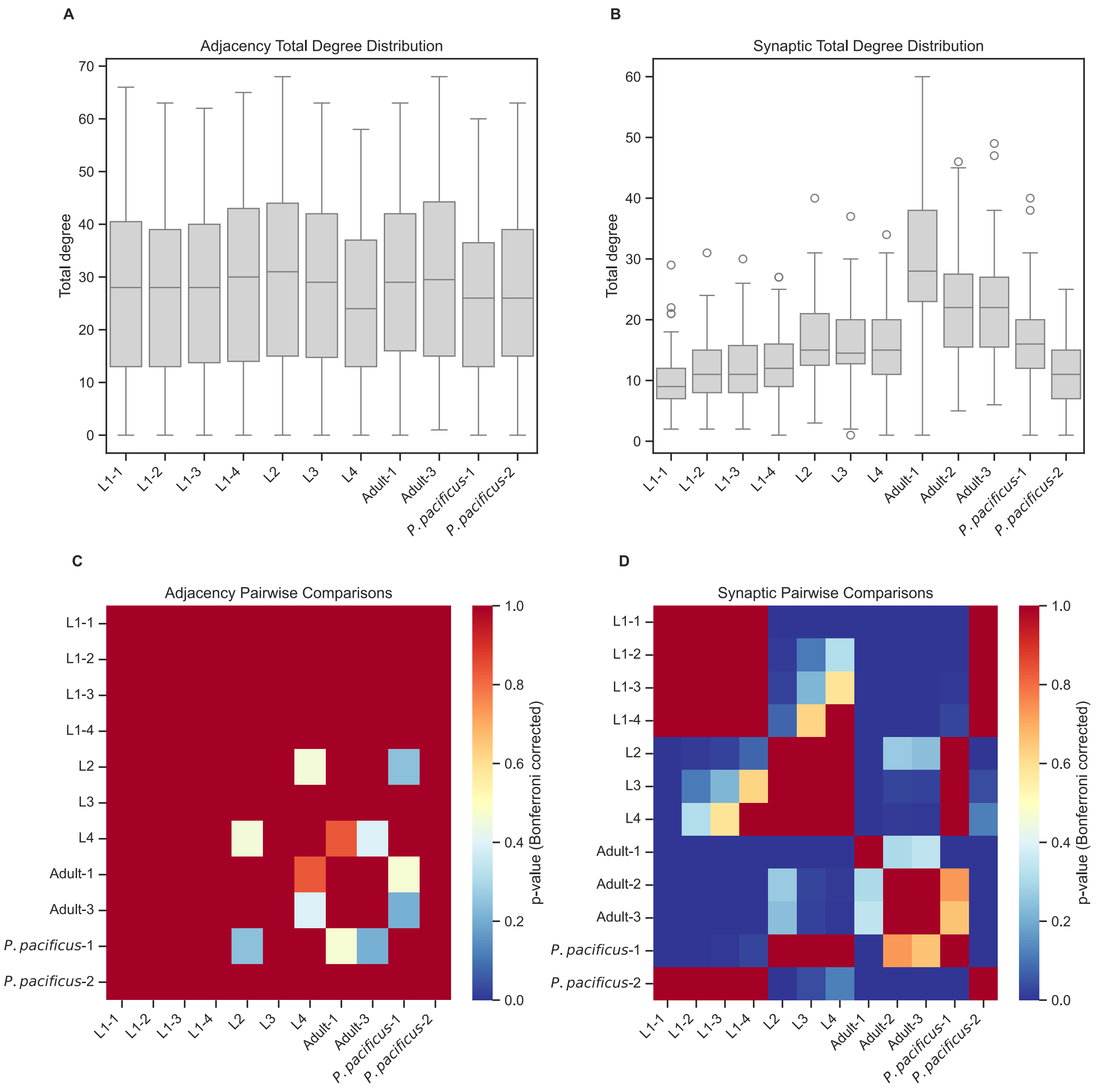


**Figure 4 - SUPPLEMENT 4: Degree distributions compared across datasets.**

**A:** Boxplots showing the distribution of total node degrees in adjacency networks across developmental stages in *C. elegans* (L1 through Adult) and adult *P. pacificus*.

**B:** Boxplots showing the distribution of total node degrees in synaptic networks across developmental stages in C. elegans (L1 through Adult) and adult P. pacificus.

**C:** Statistical comparison of adjacency network degree distributions using Dunn's test with Bonferroni correction. The heatmap shows pairwise p-values between all developmental stages, with darker colors indicating lower p-values.

**D:** Statistical comparison of synaptic network degree distributions using Dunn's test with Bonferroni correction. The heatmap shows pairwise p-values between all developmental stages, with darker colors indicating lower p-values. P-values are corrected for multiple comparisons using the Bonferroni method. The heatmap shows pairwise p-values between all developmental stages.

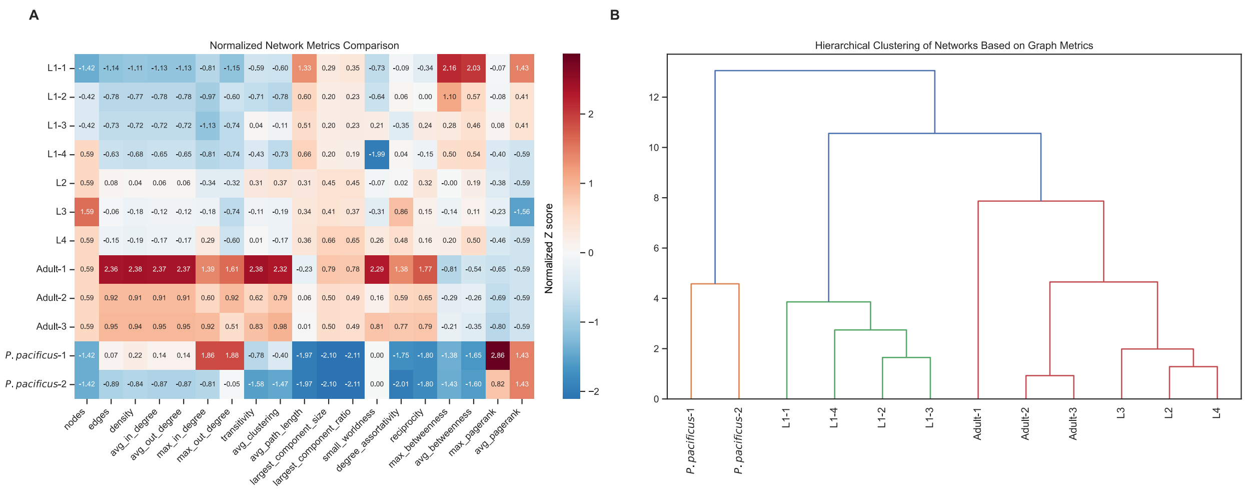


**Figure 4 - SUPPLEMENT 5: Comparative analysis of network architecture across datasets and species.**

**A:** Heatmap showing normalized network metrics for synaptic connectomes across *C. elegans* development and *P. pacificus*. Each row represents a connectome dataset, and each column represents a different network metric. Values are z-score normalized (mean=0, standard deviation=1). Blue indicates values below the mean (negative z-scores), white indicates values near the mean (z-scores close to 0), and red indicates values above the mean (positive z-scores). Network metrics include Density: Proportion of possible connections that are present, Average/Maximum In/Out Degree: Mean and maximum number of incoming/outgoing connections per neuron, Transitivity: Probability that adjacent nodes are connected (clustering), Average Clustering: Mean clustering coefficient across all nodes, Average Path Length: Mean shortest path length between all node pairs, Small-worldness: Ratio of clustering to path length relative to random networks, Degree Assortativity: Correlation between degrees of connected nodes, Reciprocity: Proportion of bidirectional connections, Maximum/Average Betweenness: Measures of node centrality in information flow, Maximum/Average PageRank: Measures of node importance in the network

**B:** Hierarchical clustering dendrogram showing relationships between connectomes based on their network metrics. The vertical axis represents the distance (dissimilarity) between clusters, with greater heights indicating greater differences. Clustering was performed using Ward's minimum variance method. The height of each branch in the dendrogram represents the increase in within-cluster variance resulting from the merger of two clusters. Shorter heights indicate more similar network properties between the merged groups.

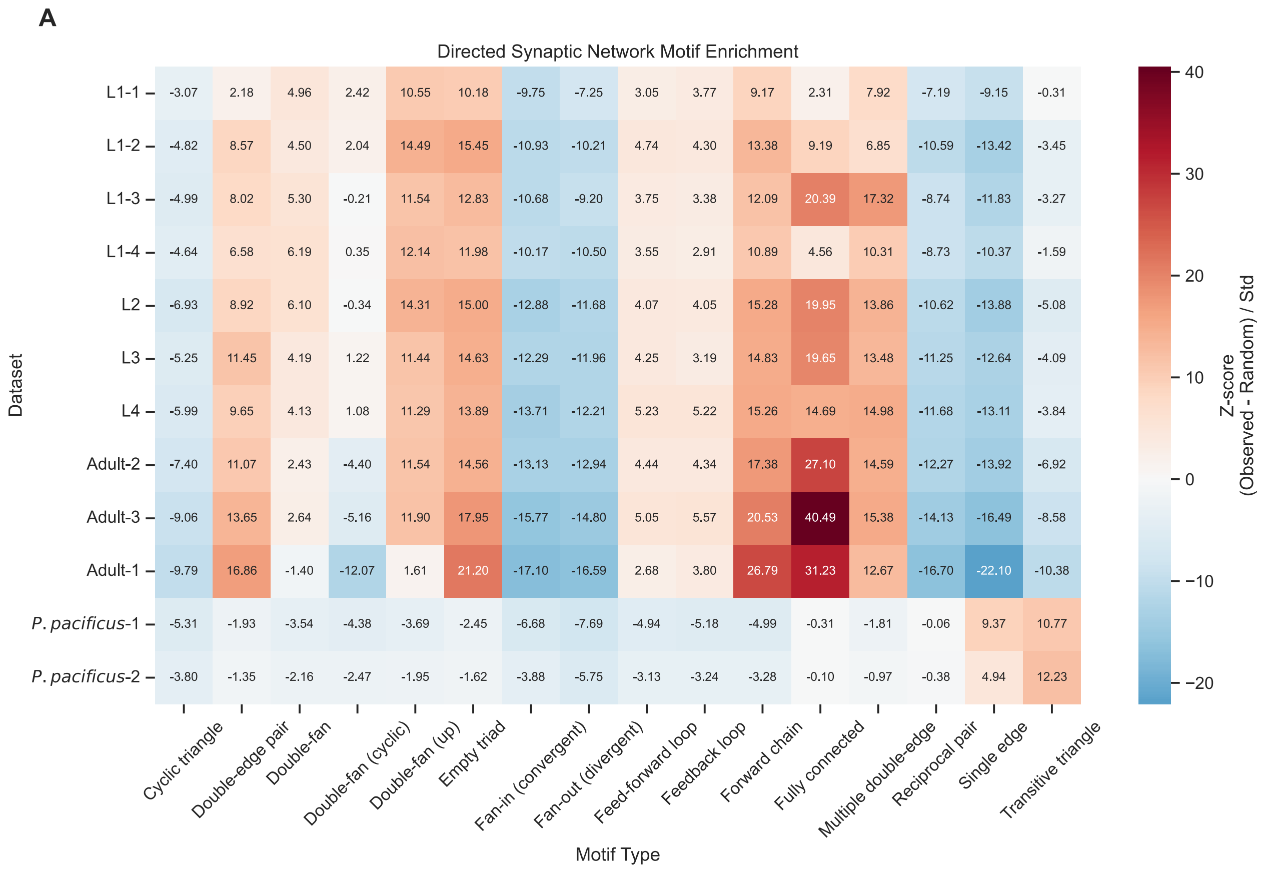


**Figure 4 - SUPPLEMENT 6: Network motif analysis reveals species-specific circuit architecture.**

**A:** Enrichment analysis of directed motifs in synaptic networks across *C. elegans* development and *P. pacificus*. Motif types include Empty triad: Three unconnected nodes, Single edge: One directed connection among three nodes, Forward chain: Sequential A→B→C connections, Fan-in: Convergent inputs (two nodes projecting to one), Fan-out: Divergent outputs (one node projecting to two), Reciprocal pair: Bidirectional connection with third unconnected node, Feed-forward loop: A→B→C with direct A→C connection, Feedback loop: Circular A→B→C→A pattern, Transitive triangle: Feed-forward pattern with all possible forward connections, Cyclic triangle: Bidirectional connections between all nodes, Double-edge pair: Two reciprocal connections, Double-fan: Complex patterns with multiple reciprocal connections, Fully connected: All possible connections present. For A and B, the heatmap shows z-scores comparing observed motif frequencies to those in degree-preserved randomized networks. Red indicates enrichment (positive z-scores) and blue indicates depletion (negative z-scores) relative to random networks. Z-scores were calculated by comparing observed motif counts to distributions from 100 randomized networks that preserved each node's degree sequence.

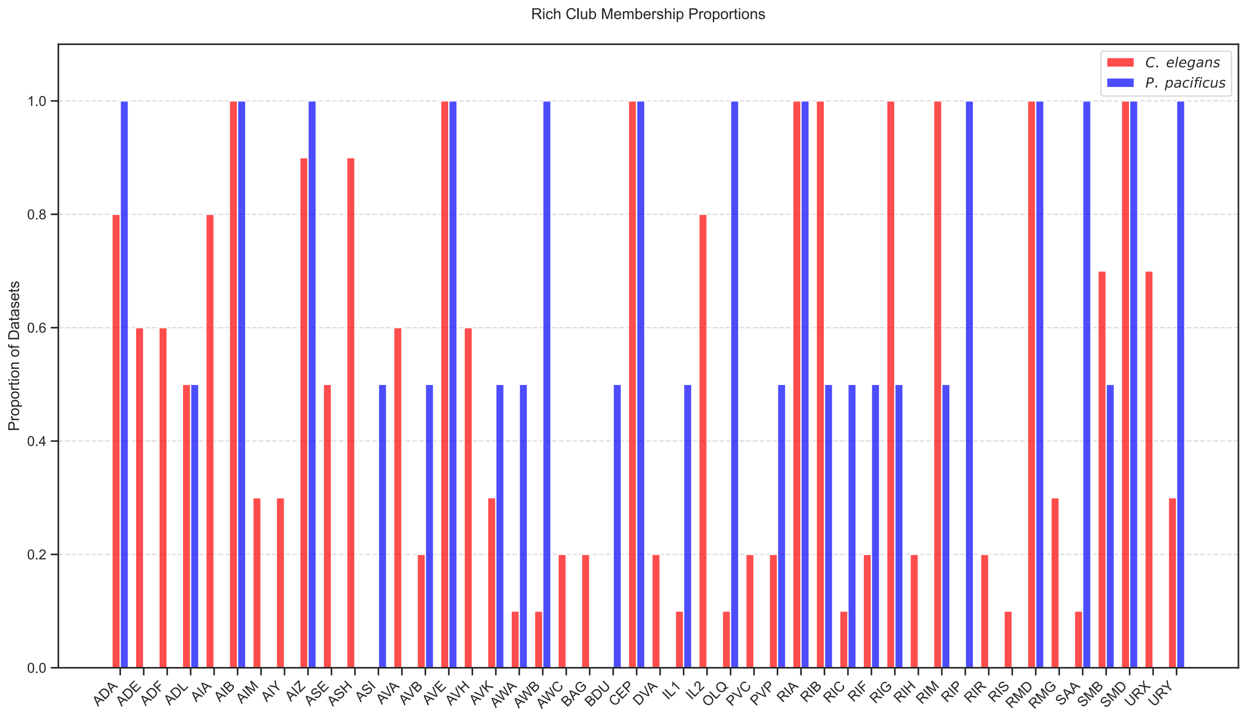


**Figure 4 - SUPPLEMENT 7: Rich club analysis across species.**

Bar plot showing the proportion of datasets in which each neuron participates in the rich club structure across *C. elegans* (red) and *P. pacificus* (blue) connectomes. Rich club neurons were identified using stringent criteria to ensure robust identification of core circuit components including Statistical Significance: Rich club coefficient ≥3 standard deviations above random expectation, based on 1000 degree-preserved randomized networks, Topological Robustness: Enrichment must occur across ≥3 consecutive degree thresholds, Size Requirements: Rich clubs must contain ≥5 neurons, Consistency Threshold: Neurons must appear in rich clubs of ≥30% of datasets. The x-axis shows individual neurons that met these conservative criteria in at least one species. The y-axis represents the proportion of datasets (0-1) in which each neuron was identified as a rich club member. Rich club analysis was performed by first calculating the rich club coefficient φ(k) for each degree threshold k, defined as the density of connections between nodes with degree ≥k, computed across all possible k values up to the maximum degree in each network. To assess statistical significance, we generated 1000 randomized networks that preserved both the in- and out-degree sequence of each node while maintaining network size and density.
